# Distinct prelimbic cortex ensembles encode response execution and inhibition

**DOI:** 10.1101/2025.02.23.639736

**Authors:** Rajtarun Madangopal, Yuan Zhao, Conor Heins, Jingfeng Zhou, Bo Liang, Giovanni Barbera, Ka Chun Lam, Lauren E. Komer, Sophia J Weber, Drake J Thompson, Yugantar Gera, Diana Q Pham, Katherine E Savell, Brandon L Warren, Daniele Caprioli, Marco Venniro, Jennifer M. Bossert, Leslie A Ramsey, Hank P. Jedema, Geoffrey Schoenbaum, Da-Ting Lin, Yavin Shaham, Francisco Pereira, Bruce T. Hope

**Affiliations:** Behavioral Neuroscience Research Branch, Intramural Research Program, National Institute on Drug Abuse, Baltimore, MD, USA; Machine Learning Core, Intramural Research Program, National Institute of Mental Health, Bethesda, MD, USA; Cellular and Neurocomputational Systems Branch, Intramural Research Program, National Institute on Drug Abuse, Baltimore, MD, USA

## Abstract

Learning when to initiate or withhold actions is essential for survival and requires integration of past experiences with new information to adapt to changing environments. While stable prelimbic cortex (PL) ensembles have been identified during reward learning, it remains unclear how they adapt when contingencies shift. Does the same ensemble adjust its activity to support behavioral suppression upon reward omission, or is a distinct ensemble recruited for this new learning? We used single-cell calcium imaging to longitudinally track PL neurons in rats across operant food reward Training, Extinction and Reinstatement, trained rat-specific decoders to predict trial-wise behavior, and implemented an *in-silico* deletion approach to characterize ensemble contributions to behavior. We show that operant training and extinction recruit distinct PL ensembles that encode response execution and inhibition, and that both ensembles are re-engaged and maintain their roles during Reinstatement. These findings highlight ensemble-based encoding of multiple learned associations within a region, with selective ensemble recruitment supporting behavioral flexibility under changing contingencies.

## Introduction

Learned stimulus–response–outcome sequences are central to survival and can guide response execution (or inhibition) in pursuit (or avoidance) of the original outcome. The medial prefrontal cortex (mPFC) plays a crucial role in various aspects of learned appetitive and aversive behaviors, such as attention, decision-making, and memory (*1–6*). Early inactivation studies using the extinction-reinstatement model led to the hypothesis that the dorsal (prelimbic cortex, PL) and ventral (infralimbic cortex, IL) mPFC subdivisions play opposite roles in behavioral expression and inhibition, respectively (*7–10*). However, subsequent studies have questioned this PL-go/IL-stop dichotomy and suggest instead that sparse and intermingled populations of neurons (neuronal ensembles) within these regions (*11–20*) control the selection of distinct task-dependent responses, rather than merely promoting or inhibiting behavior (*6, 21–23*).

In line with this hypothesis, *in vivo* electrophysiological recordings have identified diverse cue, response/inhibition, and reward-related response patterns at the single-neuron level in both subregions across multiple behavioral tasks (*21, 22, 24, 25*). However, as most studies used electrode arrays or bundles, they were limited to sparse sampling within the region during a given session and were not optimized to identify concerted population-level activity patterns (i.e., ensembles) during behavior. More recent studies applied single-cell resolution calcium imaging to simultaneously monitor hundreds of PL neurons during behavior (*26–29*) and identified PL ensembles mapping onto behavioral sequences during operant food self-administration training (*29*), as well as unique PL ensembles during Pavlovian reward learning (*26*). Interestingly, these ensembles showed stable coding patterns established during learning and could be used to infer task states (cue/trial type, reward delivery) and predict both task-related (press/lick) and non-task-related behaviors during the imaging session.

What happens to these stable ‘training’ ensembles when individuals adjust their behavior in response to changing reward contingencies, such as suppressing responses during extinction (reward omission) or re-engaging in reward-seeking during reinstatement? Does behavioral flexibility rely on changes within the same ensemble’s activity patterns, or does it involve the recruitment of distinct PL neuron populations tailored to each new behavioral context? Specifically, we asked whether the same PL ‘training’ ensemble supports training, extinction, and reinstatement of responding for a reward through changing activity patterns, or if separate non-overlapping ensembles are recruited to mediate the appropriate action during each of these phases. To answer this question, we used miniature epifluorescent microscopes (*28, 30, 31*) for longitudinal single-cell resolution calcium imaging of PL neurons in rats first as they made operant responses to receive palatable food pellet rewards in a trial-based procedure (Training), next as they learned to suppress responding when the reward was omitted (Extinction), and finally when they reinstated responding upon receiving a non-contingent ‘priming’ pellet (Reinstatement) a manipulation known to reinstate operant responding after extinction (*32, 33*). We analyzed *active neuron populations and activity patterns* as rats modified their behavior and developed decoders operating on individual rats’ PL activity and *in silico* deletion tests to determine whether the same PL ensemble neurons support training, extinction, and reinstatement of responding for food reward through changing activity patterns, or if separate non-overlapping ensembles are recruited to mediate the appropriate action during each of these phases.

## Results

### Training, extinction, and reinstatement of palatable food seeking using a trial-based operant procedure

Prior to imaging, we trained rats to self-administer palatable food pellets (*32, 34*) in three stages consisting of increasing numbers of trials and progressively shorter lever availability periods (Supplementary Figure S1B). Rats learned to make responses on the active (food-paired) lever to receive palatable food pellets during initial self-administration training (Supplementary Figure S1C, left; *Lever x Session*: F_1.8,14.6_ = 12.85, *p* < 0.001), and maintained stable responding during both phases of post-surgery training (Supplementary Figure S1C, center and right; *Lever x Session*: phase 1: F_3.2,25.6_ = 9.8, *p* < 0.001; phase 2: F_4.1,32.8_ = 4.93, *p* = 0.003). Following training, we recorded activity dynamics of PL neurons during the last two sessions of food self-administration (Training sessions; T1, T2), four consecutive sessions of extinction (Extinction sessions; E1-E4), and one session of palatable food-primed reinstatement (Reinstatement session; R) (Figure 1B, 100 trials / session). Rats maintained stable lever pressing during the two imaged self-administration training sessions, decreased lever pressing across the four imaged extinction sessions, and reinstated lever pressing in response to non-contingent ‘priming’ food pellet delivery during reinstatement testing (*Lever x Session*: F_1.3,9.02_ = 18.75, *p* = 0.001). Detailed statistical results for behavior during training and imaging sessions are provided in Supplementary Table S1.

**Figure 1.**
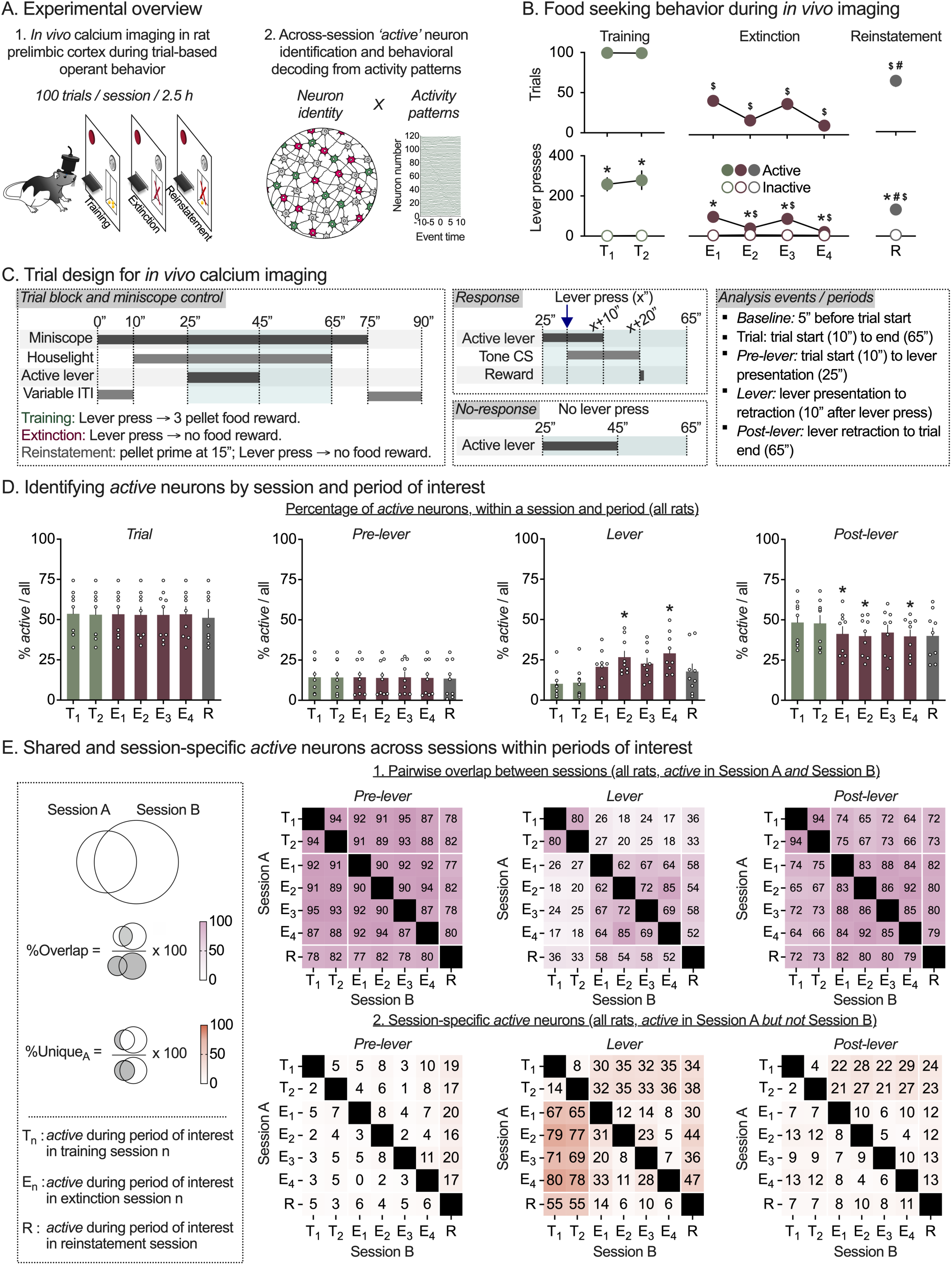
*In vivo* calcium imaging allows longitudinal tracking of neuronal activity patterns and *active* neuron overlap in rat prelimbic cortex during palatable food-seeking behavior. **(A)** *Experimental overview.* Rats were trained to self-administer palatable food pellets using a trial-based operant procedure prior to longitudinal *in vivo* calcium imaging during palatable food self-administration (Training (T), 2 sessions), extinction of palatable food seeking (Extinction (E), 4 sessions), and reinstatement of palatable food-seeking behavior by presentation of a priming pellet (Reinstatement (R), 1 sessions). Neuron spatial and temporal footprints were detected using imaging data concatenated across all 7 imaging sessions. (See supplementary figure S1 for detailed experimental timelines, behavioral training prior to imaging sessions, GRIN lens placements, and example imaging field of view). **(B)** *Palatable food seeking behavior during in vivo calcium imaging sessions.* Mean (± SEM) number of response trials (top row), and active and inactive lever presses (bottom row) during training (2 sessions, left panel), extinction (4 sessions, center panel) and pellet-primed reinstatement (1 session, right panel). *Significant difference (*p* < 0.05) between active and inactive lever presses. ^$^Significant decrease (*p* < 0.05), versus training session 1 in number of response trials (top) and active lever presses (bottom). ^#^Significant increase (*p* < 0.05), versus extinction session 4 in number of response trials (top) and active lever presses (bottom) during reinstatement. **(C)** *Trial design and analyzed events.* Temporal sequence of events under experimenter control (left), or dependent on rats’ behavioral response (middle, top) or no-response (middle, bottom). Description of trial periods analyzed (right). **(D)** *Percentage of active neurons by trial period and imaging session.* For each rat, spikes were estimated from calcium transients and used to calculate trial-by-trial average firing rate of individual neurons (all neurons detected across all 7 sessions) within each trial period. For each session and trial period, *active* neurons were identified as those with significantly different average firing rates during that period vs. baseline period at the start of the same trial in the session. Mean (± SEM) percentage of *active* neurons identified during the entire trial (left), pre-lever period (center left), lever availability period (center right) and post-lever period (right), for each imaging session. Clear circles represent data from individual rats (*n* = 9). *Significant difference (*p* < 0.05) in percentage of *active* neurons vs. Training session 1. (See supplementary Figure S2 for *active* neuron counts for each session and period, and supplementary Figure S3 for *active* neuron counts by session type.). (E) *Shared and session-specific active neurons across sessions within trial periods.* Percentage pairwise shared (1) and session-specific (2) *active* neurons were calculated for each rat, and trial period as shown in the schematic (left). Heatmaps show mean pairwise %overlap (1) and %unique (2) *active* neurons for pre-lever period (left), lever period (middle) and post-lever period (right), for all 7 imaging sessions. (See supplementary Figure S3 for heatmaps of %overlap and %unique neurons for pair-wise comparisons between 3 session types.)

Following imaging, we applied image processing routines to extract fluorescence time-courses and spatial locations of neurons for each rat using concatenated imaging data from all 7 imaging sessions (Min/Median/Max = 124/159/350 neurons, *n* = 9 rats). We used individual rats’ PL neuronal activity patterns and trial-wise behavioral measures to: (1) quantify the number and percentage of neurons active during specific session types and trial periods; (2) analyze the proportions of overlapping and unique neurons between session types and trial periods; (3) train rat-specific decoders to predict trial-wise behavioral responses using PL neuronal activity during Training and Extinction sections, and (4) test the contributions of specific neuron populations of interest to behavioral decoding accuracy during operant Training, Extinction, and Reinstatement.

### Number and percentage of active neurons by session and trial period

We first broadly quantified PL activity levels during different stages of behavior. For each rat, we estimated neuron spiking from calcium transients and identified *active* neurons during four periods of interest (Figure 1C) – (1) <u>trial period</u>: start to end of behavioral trial, (2) <u>pre-lever period</u>: trial start to active lever presentation, (3) <u>lever period</u>: active lever presentation to retraction, and (4) <u>post-lever period</u>: active lever retraction to trial end. For each session, a neuron was deemed *active* in a period if its firing rate within that period was significantly different from that of the baseline period 5 seconds prior to start of the trial (Figure S2A). For each rat, we divided the *active neuron* counts (Figure S2B) by the total number of detected neurons and compared *percent active* neurons within each trial period, across the 7 imaging sessions (Figure 1D), or collapsed by session-type (3 session types: Training (T), Extinction (E), and Reinstatement (R); Figure S3A). Overall, the percentage of active neurons increased during the lever availability period and decreased during the post-lever period during Extinction sessions (relative to Training and Reinstatement sessions), while the percentage of active neurons was unaltered for the pre-lever period and for the entire Trial period across sessions.

The repeated-measures 2-way ANOVA of *Percent active neurons* across all 7 sessions, which included within-subjects factors of Imaging session (7 sessions: T1-2, E1-4, and R; Figure 1D), and Trial period (pre-lever, lever or post-lever), showed a significant interaction between the two factors (F_2.7,21.9_ = 15.63, *p* < 0.001). Follow-up 1-way ANOVAs were significant for the lever period (F_2.3,18.6_ = 13.16, *p* < 0.001) and post-lever period (F_2.8,22.7_ = 11.38, *p* < 0.001). Post-hoc analysis indicated a significant increase in percent active lever period neurons (E2, E4 vs. T1) and a significant reduction in percent active post-lever period neurons (E1, E2, and E4, vs. T1) during extinction imaging sessions. In contrast, similar levels of activity were observed across sessions when considering the entire trial period (F_1.9,15.5_ = 1.89, *p* = 0.19), or the pre-lever period (F_1.9,14.9_ = 1.01, *p* = 0.38). The same pattern was maintained when we collapsed the sessions by type (3 session types: T, E, R; Supplementary Figure S3A). Detailed statistical results are provided in Supplementary Tables S2 (7 sessions) and S3 (3 session types).

### Overlapping versus unique active neurons by session and trial period

Next we asked whether the same (overlapping) or different (unique) sets of neurons are active over the 7 imaged sessions. For each rat and period of interest, we identified overlapping versus unique active neuron populations between pairs of the 7 imaged sessions (Figure 1E), and between pairs of the 3 session-types (Supplementary Figure S3). We calculated (1) Overlapping neurons - the percent of *active* neurons shared between two sessions/session-types as %Overlap = active in both Sessions A and B / active in Sessions A or B; i.e. |A∩B| / |A∪B| (Figure 1E.1, Supplementary Figure S3B) and (2) Unique neurons - the percent of *active* neurons that were unique to a session/session-type as %Unique_A_ = active in Session A but not Session B / active in Session A = |A-B| / |A| (Figure 1E.2, Supplementary Figure S3C).

**Figure 2.**
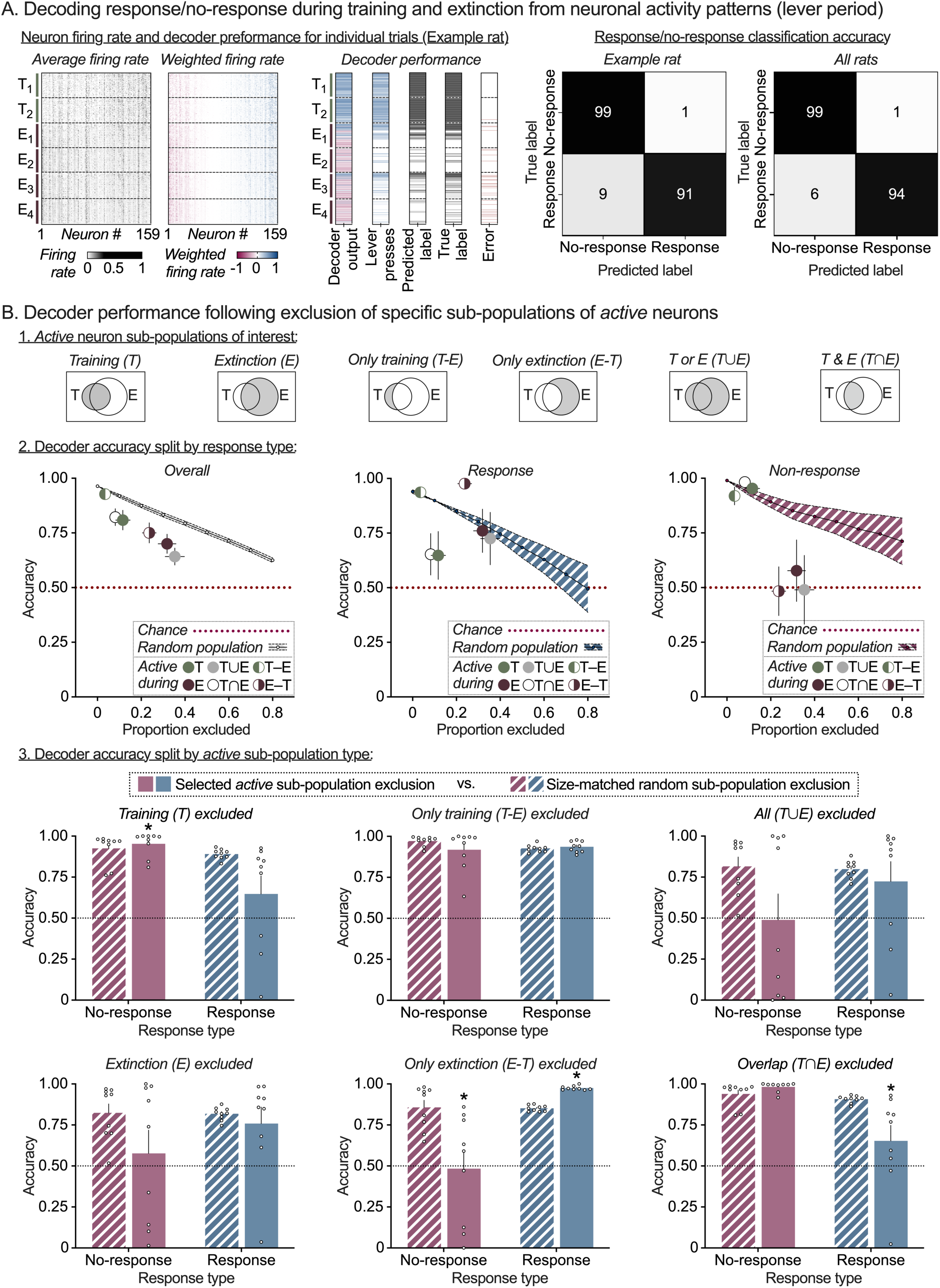
Distinct prelimbic cortex activity patterns support response/no-response behavior during Training and Extinction sessions. **(A)** *Prelimbic cortex neuronal activity supports trial-by-trial behavioral decoding during Training and Extinction.* For each rat, a binary (response/no-response) linear decoder was trained using the average firing rate vector during lever availability period, for a randomly selected subset of training and extinction trials (75/25 stratified split between training and test trials). The decoder was then tested on the holdout subset of training and extinction trials and showed high decoding accuracy for all rats. Heatmaps for one example rat showing raw firing rate of each neuron (left), firing rate multiplied by the corresponding decoder weight (center) and decoder performance (right) during the 2 training and 4 extinction sessions. Each grey pixel in the left plot represents the lever-period-average firing rate of one neuron (column) in one trial (row). Each colored pixel in the center plot represents the weighted firing rate (decoder weight x firing rate) of one neuron (column) in one trial (row). Sessions and trials are displayed in chronological order from top to bottom and neurons are sorted by weight from negative (red, left) to positive (blue, right). Decoder output column shows the sum of weighted firing rates across all neurons, which determines the decoder prediction (blue = response, red = no response). Lever press column shows number of responses by trial. Predicted label column shows binarized decoder output by trial (black tick = response predicted). True label column shows binarized response pattern by trial (black tick = response). Error column shows disagreement between predicted and true labels (red tick = error). For each rat (*n* = 9), an equal number of response and no-response trials were sampled to set chance accuracy (red dotted line) at 0.5. Decoder accuracy (%true label) on the holdout test set (training and extinction) split by response and shown as a confusion matrix for one example rat (left heatmap) or averaged across all rats (right heatmap, *n* = 9). **(B)** *Distinct sub-sets of active Training and Extinction session neurons support response and no-response prediction accuracy.* For each rat, lever period decoder accuracy was measured following exclusion of one of six combinations of Training and Extinction session-specific *active* neuron populations (top row). Note that the decoder was trained prior to the exclusion; the weights of excluded neurons are set to 0. Summary plots (row two, all rats) of Mean (± SEM) overall accuracy (left), response prediction accuracy (center), and no-response prediction accuracy (right) following exclusion of each *active* population (colored circles) vs. random population exclusion (line plot, 0-80% of all neurons per rat). Bar charts (third and fourth rows, split by population) of Mean (± SEM) response (blue) and no-response (red) prediction accuracy after exclusion of *active* population of interest (solid bars), versus exclusion of a random population of the same size for each rat (patterned bars). Clear circles represent data from individual rats (*n* = 9). *Significant difference (*p* < 0.05) in response or no-response prediction accuracy between specific population exclusion and size-matched random population exclusion.

#### Overlapping versus unique neurons within and across Training and Extinction sessions

Overall, the percentage of overlapping neurons was high within pairs of Training and Extinction sessions, but not across the two session types, only during the lever period and not the pre-lever and post-lever periods. Further, the percentage of unique neurons was significantly increased in the Extinction (versus Training) session during the lever period. This suggests that Extinction session *active* lever period (but not pre-lever or post-lever period) neurons comprise two distinct populations (1) reactivated Training session *active* lever period neurons (i.e. neurons *active* during both Training and Extinction sessions) and (2) a new population of Extinction session-specific *active* lever period neurons (i.e. neurons *active* during Extinction sessions but not previously *active* during Training).

For the pre-lever period, we observed similar levels of high %Overlap within and across training and extinction session pairs (80-95%; Figure 1E.1, left; Supplementary Figure S3B, left), and a low proportion of %Unique_A_ *active* neurons during this period, irrespective of session (Figure 1E.2, left) or session-type (Supplementary Figure S3C, left). For the lever period, we observed higher %Overlap within Training (80% for T1-2) and Extinction (> 60% for E1-4) sessions, lower %Overlap across the session types (21%; Figure 1E.1, middle; Supplementary Figure S3B, middle), and a higher proportions of %Unique_A_ *active* neurons when comparing pairs of sessions belonging to different session-types (e.g. T vs. E sessions) compared to within session-type pairs (e.g. T1 vs. T2, or E1 vs. E2-4; Figure 1E.2, middle; Supplementary Figure S3C, middle). Finally, for the post-lever period, we observed slightly higher %Overlap within Training and Extinction sessions compared to across the session types (Figure 1E.1, right; Supplementary Figure S3B, right) and a low proportion of %Unique_A_ *active* neurons during this period, irrespective of session (Figure 1E.2, right) or session-type (Supplementary Figure S3C, right).

#### Overlapping versus unique neurons in Reinstatement versus Training or Extinction sessions

Overall, the data suggest that many of the lever period neurons activated during *both* Training and Extinction sessions were re-activated during Reinstatement.

For the pre-lever period, we observed similar levels of %Overlap with the Reinstatement session across all training and extinction sessions (∼80%; Figure 1E.1, left; Supplementary Figure S3B, left). For the lever period, we found higher %Overlap between Reinstatement and Extinction (∼55%), compared to Reinstatement and Training (∼35%) sessions, and overall lower %Overlap of either session type with Reinstatement compared to within Training and Extinction session-types (60-80%; Figure 1E.1, middle; Supplementary Figure S3B, middle). Finally, for the post-lever period, we observed similar %Overlap when comparing Reinstatement to either Training or Extinction sessions (Figure 1E.1, right) or session-types (Supplementary Figure S3B, right). For the %Unique_A_ *active* neuron measure, we found higher %Unique_A_ *active* Reinstatement session neurons during the lever period vs pre- or post-lever periods, when comparing to Training (∼55%) but not Extinction (∼10%) sessions (Figure 1E.2, middle) or session-types (51% vs. 2%; Supplementary Figure S3C, middle).

#### Overlapping versus unique neurons across trial periods

Finally, we investigated the relationship between *active* neurons identified during the three trial periods. We calculated %Overlap (Supplementary Figure S4A) between active neurons identified in the three periods within each session and saw low across-period overlap and for all imaging sessions. Further, the %Unique_A_ (Supplementary Figure S4B) lever period active neurons (compared to the other two periods), stayed consistent across the 7 imaging sessions, suggesting a stable ensemble representation during the lever availability period. In contrast, for the pre and post-lever periods, %Unique_A_ neurons, were similar within session-types (i.e., T1-T2, or E1-E4), but differed across session types, likely reflecting different outcomes / behavioral patterns in these periods (e.g. pellet delivery, food port entry) between the session types.

### Decoding trial-wise behavioral responses from neuronal activity

In the analyses above, we observed stable *active* neuron populations within Training and Extinction session types, and re-activation of many of these neurons during the Reinstatement session. These patterns were specific to the lever period, suggesting the possibility that activity patterns of PL neurons during this period might contain information about rats’ operant response (or omission thereof) on the presented lever. To determine whether this is the case, we implemented rat-specific decoders to predict their response execution and inhibition behaviors following training and extinction learning, based on neuronal activity patterns. For each rat, we trained a binary linear decoder to predict trial-wise behavioral outcomes (response/no-response) from the vector of average firing rates of all neurons. We assessed two periods of interest - lever period and pre-lever period. Within each period, we trained rat-specific decoders using a randomly selected subset of the data (75% of training and extinction trials) and tested the decoder on two test sets: (1) the 25% of training and extinction trials held out from decoder training (Figure 2), and (2) 100% of trials in the reinstatement session (Figure 3). Test set 1 evaluates whether the decoders generalize to unseen training/extinction trials. Test set 2 evaluates whether the decoders work in a new context -- reinstatement session -- and, therefore, whether the information they extract from the neural activity pattern is still present then. For each test set, we sampled equal numbers of response and no-response trials to ensure 50% chance level (i.e. chance accuracy = 0.5).

**Figure 3.**
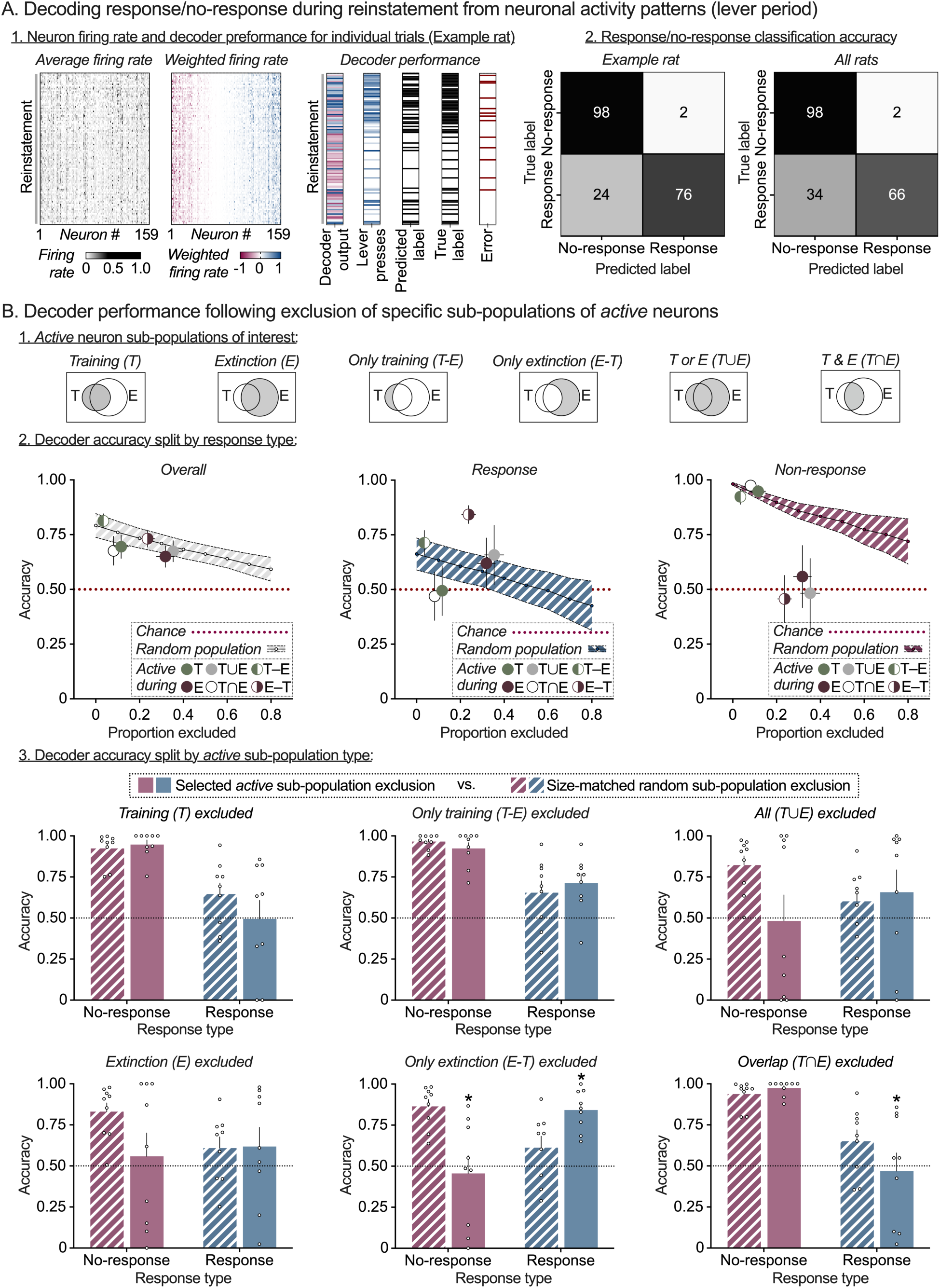
Prelimbic cortex Training and Extinction neuron activity patterns support response/no-response behavior during pellet-primed reinstatement. **(A)** *Prelimbic cortex neuronal activity supports trial-by-trial behavioral decoding during Reinstatement.* For each rat, the response/no-response decoder trained using training and extinction session trials was tested on the reinstatement session trials. Heatmaps for one example rat showing raw firing rate of each neuron (left), firing rate multiplied by the corresponding decoder weight (center) and decoder performance (right) during the reinstatement session. Each grey pixel in the left plot represents the lever-period-average firing rate of one neuron (column) in one trial (row). Each colored pixel in the center plot represents the weighted firing rate (decoder weight x firing rate) of one neuron (column) in one trial (row). Trials are displayed in chronological order from top to bottom, and neurons are sorted by weight from negative (red, left) to positive (blue, right). Decoder output column shows the sum of weighted firing rates across all neurons, which determines the decoder prediction (blue = response, red = no response). Lever press column shows number of responses by trial. Predicted label column shows binarized decoder output by trial (black tick = response predicted). True label column shows binarized response pattern by trial (black tick = response). Error column shows disagreement between predicted and true labels (red tick = error). For each rat (*n* = 9), equal number of response and no-response trials were sampled from the reinstatement session to set chance accuracy at 0.5. Decoder accuracy (%true label) on reinstatement trials split by response and shown as a confusion matrix for one example rat (left heatmap) or averaged across all rats (right heatmap, *n* = 9). **(B)** *Distinct sub-sets of active Training and Extinction session neurons support response and no-response prediction during Reinstatement.* For each rat, lever period decoder accuracy was measured following exclusion of one of six combinations of Training and Extinction session-specific *active* neuron populations (top row). Summary plots (row two, all rats) of Mean (± SEM) overall accuracy (left), response prediction accuracy (center), and no-response prediction accuracy (right) following exclusion of each *active* population (colored circles) vs. random population exclusion (line plot, 0-80% of all neurons per rat). Bar charts (third and fourth rows, split by population) of Mean (± SEM) response (blue) and no-response (red) prediction accuracy after exclusion of *active* population of interest (solid bars), versus exclusion of a random population of the same size for each rat (patterned bars). Clear circles represent data from individual rats (*n* = 9). *Significant difference (*p* < 0.05) in response or no-response prediction accuracy between specific population exclusion and size-matched random population exclusion.

A linear decoder uses the average firing rate of each neuron during the lever period in each trial as input (heatmap shown in Figure 2A, left). The decoder computes a weighted linear combination of the firing rates of all neurons (firing rates multiplied by the corresponding decoder weights are shown in Figure 2A, center left) to predict whether or not the rat will press the active lever: the prediction is yes if the linear combination is positive, or no if the linear combination is negative. Figure 2A, center right, shows the linear combination over every trial in the 2 training and 4 extinction sessions in the first column. The adjacent columns show, respectively, whether the lever was pressed in each trial, the decoder prediction, the true label (same as lever press), and the trials where the decoder made an error. Figure 2A, right, breaks down decoder performance as a confusion matrix (% of trials where prediction matches the true response, or not, for each response), both for the example rat used in the rest of the figure (Figure 2A.2, left heatmap), or averaged across all rats (Figures 2A.2, right heatmap). Figure 3A provides the analogous plots for all the trials in the Reinstatement session. Individual decoder accuracy confusion matrices (split by rat and decoder training period) are provided in Supplementary Figures S5 and S6. Detailed results for decoder accuracy on the two test sets are provided in Supplementary Tables S4 and S6, respectively.

#### Trial-wise decoding during Training and Extinction

The decoder trained on lever period activity exhibited overall high accuracy (Mean: 0.96, SEM: 0.005; Cohen’s *d* = 30.5) with slightly higher accuracy for no-response prediction (Mean: 0.99, SEM: 0.001; Cohen’s *d* = 90.8) versus response prediction (Mean: 0.94, SEM: 0.008; Cohen’s *d* = 18.1). In contrast, for the decoder trained on pre-lever period activity, overall accuracy was at chance level (Mean: 0.5, SEM: 0.01; Cohen’s *d* = 0.02) and similar for no-response prediction (Mean: 0.62, SEM: 0.11; Cohen’s *d* = 0.37) and response prediction (Mean: 0.38, SEM: 0.11; Cohen’s *d* = −0.37).

#### Trial-wise decoding during Reinstatement

The decoder trained on lever period activity also had overall high accuracy during reinstatement testing (Mean: 0.79, SEM: 0.0.055; Cohen’s *d* = 1.8) with slightly higher accuracy for no-response prediction (Mean: 0.9, SEM: 0.08; Cohen’s *d* = 1.65) versus response prediction (Mean: 0.74, SEM: 0.059; Cohen’s *d* = 1.37). In contrast, for the decoder trained on pre-lever period activity, overall accuracy was at chance level (Mean: 0.54, SEM: 0.05; Cohen’s *d* = −0.38) and similar for no-response prediction (Mean: 0.6, SEM: 0.12; Cohen’s *d* = 0.37) and response prediction (Mean: 0.38, SEM: 0.1; Cohen’s *d* = 0.28).

In agreement with the previous overlapping and unique neurons matrices, these decoding analyses indicate that stable PL activity patterns are established during the lever period as the rats learn to make, and suppress, responses over the course of operant Training and Extinction learning. These patterns can be used to predict response execution or inhibition during individual trials during learning (in the Training and Extinction sessions), and the decodability is maintained when rats re-engage with the lever during the Reinstatement session. In contrast, it appears these stable patterns are not present during the pre-lever period.

### Contribution of Active Neuron Sub-populations to Response/No-response Prediction

In the previous section, we established the existence of distinct, stable patterns of activity during the lever period that are associated with response execution or inhibition, respectively. The next question we posed is which specific sub-populations of active neurons contain the information that the decoder uses in predicting response with high accuracy. For each rat, we assessed the contribution of six sub-populations of Training and Extinction session-specific *active* neurons to lever period decoder performance during training and extinction (Figure 2B) or reinstatement (Figure 3B). The six sub-populations of *active* neurons considered were (1) Training (T) – active in any of the 2 training sessions; (2) Extinction (E) - active in any of the 4 extinction sessions; (3) Only Training (T-E) – active in training but not extinction sessions; (4) Only Extinction (E-T) – active in extinction but not training sessions (i.e. the putative *Extinction* ensemble); (5) All T or E (T ∪ E) – active in either training, or extinction sessions; (6) Overlap between T & E (T ∩ E) – active in both training and extinction sessions (i.e. the putative *Training* ensemble).

We used an *in-silico lesion* approach to test the contribution of each of the six sub-populations to trial-wise response/no-response prediction accuracy of the previously trained lever period decoder. To this effect, we removed specific sub-populations from decoder prediction by setting their corresponding decoder weights to 0 when computing the linear combination of neuron firing rates and calculated the accuracy of the resulting decoder (*specific-population-exclusion accuracy*). For a comparison controlling for accuracy changes based on sub-population size, we also calculated the accuracy of a decoder obtained after exclusion of a randomly chosen size-matched sets of neurons (*random-population-exclusion accuracy*). Summary plots of overall accuracy (left), response prediction accuracy (center), and no-response prediction accuracy (right) following exclusion of each active sub-population (colored circles) vs. random population exclusion (0-80% of all neurons per rat, line with error band) are shown in Figures 2B.2 and 3B.2.

We first used 2-way repeated measures ANOVA with within-subjects’ factors of *Population* (1-6) and *Drop type* (specific, random) to test whether the sub-populations have enhanced contributions to overall decoder prediction compared to randomly selected PL neurons. Then for each sub-population, we used 2-way repeated measures ANOVA with within-subjects’ factors of *Drop type* (specific, random) and *Response type* (response, no-response) to assess whether sub-population exclusion resulted in biased effects on response vs no-response prediction accuracy (Figures 2B.3 and 3B.3). Detailed results following exclusion of specific *active* neuron sub-populations are provided in Supplementary Tables S5 and S7, respectively.

#### Prediction accuracy during Training and Extinction

Overall decoder accuracy (Response and No-response prediction combined) after excluding each of the 6 *active* neuron sub-populations was lower than size-matched random sub-population exclusion (Figure 2B.2, left). The overall 2-way ANOVA showed a significant *Population* x *Drop type* interaction (F_1.96,15.7_ = 2.55, *p* = 0.043) and post-hoc analysis revealed significant accuracy drop for 4 of the 6 *active* sub-populations tested (E, T∪E, T∩E, and E-T; see Supplementary Tables S4 for detailed statistical outputs). When assessing decoder accuracy separately for Response versus No-response types (Figures 2B.2, middle and right), 2-way ANOVA showed significant *Response type x Drop* type interaction for 3 of the 6 sub-populations tested (T, T∩E, and E-T; see Supplementary Tables S5 for detailed statistical outputs). Exclusion of neurons belonging to the putative *Training* ensemble (Overlap (T∩E) sub-population; Figure 2B.3, bottom right) decreased decoder accuracy to chance levels, specifically for response trials (*p* = 0.029). In contrast, exclusion of neurons belonging to putative Extinction ensemble (Only Extinction (E-T) sub-population; Figure 2B.3, bottom middle) decreased decoder accuracy to chance levels for no-response trials (*p* < 0.001) and showed a moderate increase in decoder accuracy for response trials (*p* < 0.001).

#### Prediction accuracy during Reinstatement

Overall decoder accuracy (Response and No-response prediction combined) after excluding each of the 6 *active* neuron sub-populations was lower than size-matched random sub-population exclusion (Figure 3B.2, left). The overall 2-way ANOVA showed a main effect of *Population* (F_1.64,13.1_=3.85, *p* = 0.006) but not *Drop type* (F_1,8_=2.7, *p* = 0.14), or interaction (F_2.04, 16.3_ =2.87, *p* = 0.085) between the factors (see Supplementary Tables S6). When assessing decoder accuracy separately for Response versus No-response types (Figures 3B.2, middle and right), the 2-way ANOVA showed significant *Response type x Drop* type interaction for 2 of the 6 sub-populations tested (T∩E, and E-T; see Supplementary Tables S7). Again, while exclusion of neurons belonging to the putative *Training* ensemble (Overlap (T∩E) sub-population; Figure 3B.3, bottom right) decreased decoder accuracy to chance levels, specifically for response trials (*p* = 0.012), exclusion of neurons belonging to putative Extinction ensemble (Only Extinction (E-T) sub-population; Figure 3B.3, bottom middle) decreased decoder accuracy to chance levels for no-response trials (*p* < 0.001) and showed a moderate increase in accuracy for response trials (*p* < 0.001).

Overall, the decoding analyses indicate that *active* lever period neurons were significantly associated with Response/No-response (compared to similar-sized randomly selected neuron populations) both during Training and Extinction sessions, and during Reinstatement. Further, the activity patterns of PL *Training* and *Extinction* ensembles supported opposite response predictions - *Training* ensemble activity supported response execution and Extinction ensemble activity supported response inhibition on individual trials, across Training, Extinction and Reinstatement.

## Discussion

We used single-cell resolution calcium imaging to monitor the activity patterns of hundreds of PL neurons in rats during trial-based operant food reward training, extinction, and reinstatement. Our goal was to test whether the same PL ensemble neurons adapt their activity patterns to support training, extinction, and reinstatement of reward-seeking behavior, or if distinct, non-overlapping ensembles are recruited to mediate the appropriate action during each phase. Population-level analyses of overlapping versus unique *activ*e neurons across the Training and Extinction sessions revealed higher overlap for lever period *active* neurons within the same session type than between them; this was not the case for the pre-lever or post-lever periods. Additionally, many *active* lever period neurons from Training were re-engaged during Extinction, and a new population of *active* neurons emerged specifically during Extinction.

We trained rat-specific decoders to predict trial-wise response/no-response behavior based on PL *activity patterns* during training and extinction. Decoders trained on lever period activity patterns accurately predicted individual rats’ trial-wise response (execution) versus no-response (inhibition) behavior; this was not the case for the pre-lever period.

Next, we tested the contribution of specific sub-populations of *active* Training and Extinction session neurons by omitting them from the decoder (i.e., an *in silico* targeted deletion. Omission of lever-period neurons *active* during both Training and Extinction sessions (T∩E) selectively reduced response prediction accuracy to chance levels, suggesting these represent a *Training ensemble*, while omission of lever-period neurons *active* only during Extinction (E-T) disrupted no-response prediction, suggesting these represent an *Extinction ensemble*.

Finally, these opposite decoder prediction impairments after *Training* or *Extinction ensemble* omission persisted during Reinstatement, indicating that both ensembles are re-engaged and maintain their respective roles. Overall, the results of our analyses indicate that operant training and extinction learning recruit distinct PL ensembles to support behavioral flexibility in response to changing reward contingencies.

Previous in vivo recording studies have demonstrated diverse PL activity patterns related to cues, response execution/inhibition, and outcome/reward (*5, 21, 24, 25, 35*), and have identified stable PL neuron populations (ensembles) during both operant and Pavlovian reward learning (*14, 26, 29, 36*). Consistent with these findings, we observed diverse PL neuron activity patterns during Training sessions, with similar numbers and high overlap of *active* neurons across sessions, regardless of trial period. However, contrary to the PL-go/IL-stop hypothesis’ prediction of reduced PL engagement during Extinction, the number of *active* neurons increased in Extinction sessions. Notably, this increase was specific to the lever availability period and there was higher overlap in lever period *active* neurons within each session type than between Training and Extinction sessions, indicating that different ensembles encode Training versus Extinction in the same PL subregion.

Furthermore, decoders trained on PL lever period activity patterns consistently predicted individual rats’ response execution and inhibition across Training and Extinction sessions, suggesting that the PL does not simply support responding during reward availability and ‘go offline’ during Extinction. These results align with previous *in vivo* electrophysiological recording studies using a discriminative stimulus (DS) procedure for sucrose (*21, 22*) and cocaine reward, which found that neuronal activity in PL and IL is not specifically locked to the DS or the action (i.e., going or stopping), but rather reflects the appropriate behavioral action based on context (e.g., responding during DS+ versus inhibiting responses following extinction training).

Surprisingly, lever period *active* neurons from the Training session remained active during the Extinction session. Instead, shared and overlapping neuron analysis between Training and Extinction sessions revealed that a majority of lever period active neurons from the Training session were *also active* during Extinction (T∩E), along with a second population *exclusively active* during Extinction (E-T). One possibility is that both populations are simultaneously active, and that the extinction only (E-T) population suppresses response execution mediated by the re-activated Training population (T∩E). However, the response-specific bias in decoder prediction accuracy following targeted omission of these two subpopulations suggests another explanation.

Omission of the T∩E subpopulation alone selectively reduced decoder *response* prediction accuracy to chance levels, while omission of the E-T subpopulation specifically disrupted *no-response* prediction accuracy. This suggests that these two populations are likely not active and in opposition but instead come online to support separate outcomes. In this perspective, the T∩E subpopulation would represent a *Training* ensemble that is formed during Training and supports response execution in pursuit of the reward. This ensemble would then be reactivated during *response trials* in the Extinction sessions when rats continue to make responses (to earn food reward based on the previous contingency) as they learn the new contingency (i.e. extinction learning). The second E-T subpopulation would then represent the new *Extinction* ensemble that is recruited as rats learn to suppress behavior on *no-response trials* during the Extinction session.

It is possible that extended extinction training might lead to a reduction in the size of the T∩E population as rats learn to make fewer responses in pursuit of reward. We used a within day extinction procedure and chose 4 extinction sessions (based on preliminary behavioral experiments) to minimize the number of recording days and maintain stable across-session recordings. While additional experiments with longer extinction periods are needed, in support of this idea, we previously found that the number of *active* IL extinction session neurons, measured using the activity marker Fos, decreases between 2 and 7 days of extinction training (*12*). Furthermore, targeted ablation (*37*) of *active* training session neurons reduced food lever responding, while ablation of the *active* extinction session neurons increased lever responding (*12*). Another question for future research is the cell type composition of these Training and Extinction ensembles. While some studies suggest that extinction learning is mediated by preferential engagement of inhibitory, GABAergic interneurons (*35, 36*), others suggest coordinated engagement of multiple PFC cell types across reward learning (*12, 19*). Overall, our data align with recent studies showing the development of stable PL ensembles (*26, 29*) during reward learning, and suggest that operant training and extinction learning recruit distinct PL ensembles whose *activity patterns* support behavioral flexibility in response to changing reward contingencies.

Finally, *Training* and *Extinction* ensembles were re-activated during pellet-primed reinstatement and maintained their decoder selectivity for response and no-response prediction. At the population level, we observed that lever period *active* neurons from both Training and Extinction session were reactivated during the reinstatement session (although to different degrees), and very few neurons were uniquely active during the same period in reinstatement. Notably, although the rat-specific decoders were never trained on reinstatement activity patterns, lever period (but not pre-lever) decoder performance was significantly above chance levels when tested on reinstatement trials. Further, omission of active neuron sub-populations increased decoder error, and omission of the *Training* (T∩E) or *Extinction* (E-T) ensembles oppositely biased decoder prediction errors towards response versus no-response, respectively. These findings suggest that behavior during reinstatement is not driven by a distinct third ensemble but involves preferential recruitment of one of two response-outcome-specific ensembles acquired during operant training and extinction. The *Training* ensemble is engaged when rats reinstate lever pressing in response to the acute non-contingent reward provided by the priming pellet, while the *Extinction* ensemble supports response inhibition as they suppress responding when lever presses fail to result in food pellet delivery.

In summary, we show that two distinct PL ensembles are recruited over the course of operant reward training and extinction to support opposite behavioral responses, and these same ensembles are re-engaged when rats reinstate operant behavior in response to a ‘priming’ pellet. Our findings provide neuronal ensemble-based evidence supporting the idea that operant extinction reflects new learning, rather than a weakening of the response-outcome association (*38, 39*). Consistent with this view, we show that the reinstatement of operant behavior after extinction relies on the re-engagement of pre-established ensembles rather than the recruitment of a distinct third ensemble. Overall, our findings challenge the *one primary role per region* premise of the PL-go/IL-stop hypothesis and demonstrate how the interplay of pre-established and newly recruited ensembles within reward-related brain regions enables behavioral flexibility in response to changing reward contingencies.

## Materials and Methods

We combined a trial-based operant behavioral procedure with *in vivo* calcium imaging to investigate PL neuron activity dynamics during different stages of palatable food taking and seeking behavior in rats. A schematic overview of the study design is shown in Figure 1A. Timelines for surgical procedures, behavioral training, and imaging sessions are provided in Supplementary Figures S1A and S1B. We first trained rats to lever press for 45 mg palatable food pellets (*32*) using a trial-based procedure. We then used miniature epifluorescent microscopes to track PL activity patterns during operant food self-administration (Training), as they learned to suppress responding when the reward was omitted (Extinction), and when they reinstated responding after receiving a non-contingent ‘priming’ pellet (Reinstatement). We kept the microscopes attached for the entire duration of the imaging study (7 sessions over 4 days) to maintain the same imaging field of view and allow for longitudinal tracking of PL neuron activity patterns. All procedures were approved by the NIDA IRP Animal Care and Use Committee and followed guidelines outlined in the Guide for the Care and Use of Laboratory Animals. A detailed description of experimental subjects, apparatus, imaging procedures, data analysis pipelines, and statistical analyses is provided below.

### Subjects

We used male Long Evans rats (Charles River, Raleigh, *n* = 24), weighing 300-350 g at the start of the experiment. We group-housed rats in the animal facility (two per cage) for one week to minimize transportation stress, and single-housed them during the experiments. For all experiments, we maintained the rats under a reverse 12:12 h light/dark cycle (lights off at 8 AM) with free access to water in their home cages. We gave rats free access to standard rat chow for 1 week following each surgery, and maintained them at 80% ad libitum feeding during all other parts of the experiment. All procedures were approved by the NIDA IRP Animal Care and Use Committee and followed the guidelines outlined in the Guide for the Care and Use of Laboratory Animals (*40*). We excluded 4 rats based on initial training performance, 10 rats based on poor imaging field of view, and one rat due to hardware failure and excessive motion artifacts. We report the number of rats included in each analysis in the corresponding figure legend. We used only male rats because the trial-based training-extinction-reinstatement procedure, along with the calcium imaging, was established and conducted between 2015 and 2017—before we began routinely including both males and females in our studies.

### Surgery

#### Intracranial virus surgery

We anesthetized rats with isoflurane prior to injecting adeno-associated viruses (AAVs) into the prelimbic cortex (unilateral, right hemisphere). We used the following stereotaxic coordinates from bregma for virus delivery: antero-posterior (AP), + 3.2 mm; medio-lateral (ML), ± 1.3 mm; dorso-ventral (DV), −3.45 mm; 10° angle with blunt ear bars and nose bar set at −3.3 mm (Model 962, David Kopf Instruments). We used Nanofil syringes (10 uL syringe and 33 g injector needle, World Precision Instruments) attached to a ultramicropump (UMP3, World Precision Instruments) and controller (Micro4, World Precision Instruments) to deliver 500 nL of virus (AAV2/1-hSyn1-GCaMP6s, 1.9E13 GC/mL) at a rate of 100 nL/min. Following surgery, we administered once daily injections of meloxicam (1 mg/kg, subcutaneous, Covetrus, formerly Butler Schein) and dexamethasone (0.2 mg/kg, subcutaneous, Covetrus, formerly Butler Schein) for three additional days post-surgery and allowed rats to recover for at least 7 days before initial behavioral training.

#### Intracranial GRIN lens implantation surgery

Following initial behavioral training, we performed intracranial surgery to implant a 1 mm diameter, 0.95 pitch graded refractive index (GRIN) lens (ILW-100-P, Go!Foton) above the prelimbic cortex (unilateral, right hemisphere). Prior to surgeries, GRIN lenses were coated with parylene-C and fibronectin using a 2-step procedure that has been shown to minimize toxicity and maximize biocompatibility (*41*). We used the following stereotaxic coordinates from bregma for GRIN lens positioning: antero-posterior (AP), + 3.2 mm; medio-lateral (ML), ± 0.7 mm; dorso-ventral (DV), −3.5 mm; 0° angle. We administered dexamethasone (0.2 mg/kg, subcutaneous, Covetrus, formerly Butler Schein) one hour prior to surgery to reduce inflammation. We anesthetized rats using isoflurane gas (5% induction, 1-3% maintenance), and used a micro drill to create a hole in the skull above the implantation site, and installed 6 presterilized jewelers screws: two anterior to bregma, two posterior to bregma, one posterior to lambda, and one on the left hemisphere adjacent to the implantation site. We then inserted a custom tapered 1 mm diameter glass needle to 0.2 mm above the final coordinates, and held it in place for 5 minutes to create a cavity for GRIN lens implantation. We withdrew the needle, slowly inserted the GRIN lens into the final coordinates, and used dental cement to attach it to the jewelers screws on the skull. We placed a custom protective cap above the GRIN lens and secured it in place using adhesive, and sutured skin closed around the headmount assembly. We administered once daily injections of meloxicam (1 mg/kg, subcutaneous, Covetrus, formerly Butler Schein) and dexamethasone (0.2 mg/kg, subcutaneous, Covetrus, formerly Butler Schein) for seven additional days post-surgery and allowed rats to recover for up to 3 weeks before continuing behavioral training. We removed the custom protective cap once every two weeks and imaged under isoflurane anesthesia (5% induction, 1-2% maintenance) using a custom epifluorescent microscope to monitor scar tissue regression and determine the appropriate time for miniature microscope optical alignment and baseplate installation.

#### Miniscope alignment and baseplate installation

Following scar tissue regression, we performed a final non-invasive procedure to align the Miniscope to the implanted GRIN lens and installed a custom threaded baseplate (modified from Thor labs 0.5" lens tube, SM05M05) for stable optical coupling during behavior. For this procedure, we first administered dexamethasone (0.2 mg/kg, subcutaneous, Covetrus, formerly Butler Schein) one hour prior to surgery to reduce inflammation. Next we anesthetized rats using isofluorane gas mounted rats onto a stereotaxic frame using blunt earbars and nose bar set at −3.3 mm. We made a skin incision along the midline to expose the previously cemented region, removed the custom protective cap above the GRIN lens, cleaned the surface with hydrogen peroxide and sterile saline, and allowed it to dry completely before the next step. We attached the Miniscope to the baseplate using a custom threaded coupler (modified from Thor labs 0.5" lens tube end cap, SM05CP1) and used a three-axis kinematic mount (Thor labs Small Kinematic V-Clamp Mount, KM100V) coupled to a stereotaxic arm to align the Miniscope + baseplate assembly with the GRIN lens. We used tissue autofluorescence and vasculature within the field of view to optimize alignment along all three axes and adjusted the baseplate to make contact with the previously applied cement layer on the skull surface. We then used dental cement to attach the baseplate to the skull and the GRIN lens assembly, installed a removable threaded cap (Thor labs 0.5" lens tube cap SM05CP2) to protect the GRIN lens, fixed it in place using three radially placed set screws and sutured skin closed around the entire assembly. We administered once daily injections of meloxicam (1 mg/kg, subcutaneous, Covetrus, formerly Butler Schein) and dexamethasone (0.2 mg/kg, subcutaneous, Covetrus, formerly Butler Schein) for three days post-surgery, and allowed rats to recover for at least 1 week.

#### Habituation using ‘dummy’ Miniscopes

Following recovery, we attached dummy Miniscopes that had the same weight and physical tethering properties of the recording Miniscopes. We tethered the rats in the box and ran them on tethered self-administration training sessions to habituate them to the setup. We continued tethered training with dummy scopes till rats showed stable food self-administration before starting the imaging sessions.

#### Miniscope attachment for imaging

One day before the first imaging session we lightly anesthetized rats using isofluorane gas, removed the threaded protective cap, and attached the Miniscope. We lowered the scope into z alignment while observing signal from the previously aligned x-y field of view. We used vascular contrast where available, and used gentle whisker stimulation to evoke activity in the prelimbic cortex and refine Miniscope alignment. We anchored the aligned Miniscope to the baseplate using three radially placed set screws, and applied dental cement at the junctions to reduce vibration and mechanical movement during behavior. We allowed the animal to recover from anesthesia, and waited at least 1 hour before moving them to the behavior chamber. Finally, we tested the Miniscope + acquisition equipment for optical alignment and stability as animals freely explored the behavior chambers. After confirming alignment, we disconnected the data acquisition cable and housed the rats in the behavior chambers for the rest of the experiment. For each rat, we kept the Miniscope position and alignment fixed for all seven imaging sessions and attached/disconnected the data acquisition cable before/after the session to minimize cable damage.

### Behavioral apparatus

We used standard Med Associates self-administration chambers (Med Associates ENV-007) for palatable food (TestDiet, USA; Catalogue # 1811155, 12.7% fat, 66.7% carbohydrate, and 20.6% protein) self-administration training and for *in vivo* calcium imaging during stages of palatable food seeking behavior. Each chamber was enclosed in a ventilated, sound-attenuating cabinet with blacked out windows and equipped with a stainless-steel grid floor and two side-walls, each with three modular operant panels. We used a red houselight (Med Associates ENV-221M with red lens cap) located at the top of the center panel on one side of the chamber to illuminate the chamber during the training and imaging sessions. Two levers served as operant manipulanda – (1) a non-retractable lever housed in the modular panel adjacent to the houselight served as the inactive lever and (2) a retractable lever housed in the modular panel on the opposite wall (next to the food receptacle) served as the active lever.

### Behavioral procedures

We trained rats on a trial-based palatable food self-administration procedure and used *in vivo* calcium imaging to monitor activity dynamics of PL neurons during the last two sessions of food self-administration (Training sessions; T1, T2), four consecutive sessions of extinction (Extinction sessions; E1-E4), and one session of palatable food primed reinstatement (Reinstatement sessing; R) behavior. The individual stages of training and imaging are described in detail below.

#### Initial self-administration training (6 days)

We first trained male rats to lever press for palatable food reward during once daily 1-hour trial-based food self-administration sessions. We gave rats the 45-mg food pellets in their home cage one day prior to the start of training and used crushed food pellets to get rats to engage with the lever during the first 2 days of training when necessary. Each session in this phase consisted of 30 discrete trials separated by a variable inter-trial interval – the start of each trial was signaled by the illumination of a house light for 10 s, following which rats were given access to a retractable lever (active lever) for 60 s. Presses on the active lever (Fixed Ratio 1 reinforcement schedule) resulted in activation of a tone conditioned stimulus (CS) for 20-s followed by the delivery of 3 palatable food pellets. The active lever was retracted 10-s after CS onset and additional responses during this 10-s period were not reinforced. Presses on the fixed inactive lever were recorded throughout the session but had no programmed consequence.

#### Post-surgery training phase 1 (8 days)

Twenty rats successfully learned the task during this initial training. We implanted these rats with GRIN lens for imaging, allowed them to recover from surgery and then started the next phase of training consisting of once daily 2 h food trial-based self-administrations sessions. During this phase they received 60 discrete trials (separated by a variable inter-trial interval) over 2 h with 60-s lever access per trial. The reinforcement schedule and timing of events post lever press was kept the same as initial training.

#### Post-surgery training phase 2 (8 days)

All rats continued to make lever responses to get food rewards during phase 1 and were moved to the next phase consisting of once daily 3 h food trial-based self-administrations sessions. During this phase they received 100 discrete trials (separated by a variable inter-trial interval) over 2.5 h with 20-s lever access per trial. The reinforcement schedule and timing of events post lever press was again kept the same as initial training.

#### Self-administration imaging sessions (denoted as Training sessions T1 and T2)

We recorded PL calcium dynamics during 2 sessions of palatable food self-administration −1 session each on days 1 and 3 of imaging timeline with a rest day on day 2. Each 2.5 h session consisted of 100 discrete trials (separated by a variable inter-trial interval) over 3 h with 20-s lever access per trial. The reinforcement schedule and timing of events post lever press was kept the same as training. The Miniscope was triggered ON via TTL pulse 10-s before the start of each trial (signaled by houselight turning ON), and turned off 10-s after the end of the trial (signaled by houselight turning OFF). Each trial therefore consisted of approximately 75 seconds of imaging data acquired at a rate of 10.49 image frames per second from a 1.1 mm^2^ field of view (FOV), at a pixel resolution of 2.75 μm per pixel, yielding a single image size of 400 x 400 pixels, and 799 frames per trial.

#### Extinction imaging sessions (denoted as Extinction sessions E1, E2, E3 and E4)

We recorded PL calcium dynamics while rats learned to extinguish food self-administration behavior over four 3 h sessions. The 1^st^ and 2^nd^ extinctions sessions were conducted on day 3, 30 min after session T2, with a 30 min break between sessions. The 3rd and 4th extinction sessions were the first 2 sessions of day 5, with a 30 min break between sessions. The session duration, number of trials, reinforcement schedule and timing of cue presentation and lever removal post lever press was kept the same as training, but palatable food reward was withheld (extinction conditions). The Miniscope recording timeline, frame rate and field of view were kept identical to the training sessions.

#### Reinstatement imaging session (denoted as Reinstatement session R)

Following 4 sessions of extinction, we recorded PL calcium dynamics while rats exhibited palatable food-seeking behavior in response to non-contingent delivery of ‘priming’ food pellets (1 session on day 5 of imaging timeline, 30 min after extinction session E4). The key change in this session was the non-contingent delivery of a single non-contingent ‘priming’ pellet 5-s after the start of each trial. The session duration, number of trials, reinforcement schedule and timing of cue presentation and lever removal post lever press was kept the same as the extinction sessions. The Miniscope recording timeline, frame rate and field of view were also unchanged.

### Analysis of behavioral data

We conducted statistical analysis on two behavioral measures - (1) the total number of trials with at least one food-paired lever press (denoted as *Trials*) and (2) the total number of active and inactive lever responses made during the session (denoted as *Lever presses*). We used repeated measures ANOVAs with Greenhouse-Geisser correction as appropriate, and followed up on statistically significant main effects or interactions with post-hoc tests as described below. Alpha (significance) level was set at 0.05, two-tailed. Because some of our models yielded multiple main effects and interactions, we report only those that are critical for data interpretation.

For trial-based palatable food self-administration training sessions prior to imaging, we performed statistical analysis on each phase (Initial training, and post-surgery training phases 1, and 2) separately. We analyzed *Trials* using 1-way ANOVA with within-subject factor of training session and *Lever presses* using 2-way ANOVA with within-subject factors of training session and lever type (active, inactive). We used Bonferroni’s multiple comparison correction for pairwise comparisons between active and inactive lever presses within a training session.

For imaged behavioral sessions (2 training sessions, 4 extinction sessions, 1 reinstatement session), we analyzed *Trials* using 1-way ANOVA with within-subject factor of session (7 sessions) and *Lever presses* using 2-way ANOVA with within-subject factors of session (7 sessions) and lever type (active, inactive). For both measures we used Bonferroni’s multiple comparison correction for pairwise comparisons between imaging sessions.

### Calcium imaging data processing

#### Image pre-processing

We applied a sequence of image processing routines to extract fluorescence time-courses and spatial locations of single neurons from the imaging data. For each rat, we concatenated image sequences from all imaged sessions (7 sessions x 100 trials/session x 799 images/trial for each rat), and applied the NoRM-Corre non-rigid motion correction procedure (*42*) to correct for affine and non-affine shifts between images. Next, we spatially filtered all frames with a Gaussian kernel of width 16.5 μm and support 44 μm (6 and 16 pixels, respectively) to enhance high-frequency features of the data on the length scale typical of cell somas. Then we passed each pixel’s fluorescence timecourse through a threshold of 5 SD of the noise baseline for that pixel. This noise variation was estimated as the integrated spectral density within the top frequency range of the Fourier decomposition of each pixel’s fluorescence timeseries. This denoising approach has been empirically found to be effective in separating the neural contribution to fluorescence variation from those reflecting various sources of pixel noise(*43, 44*).

#### Extraction of spatial and temporal footprints

After thresholding the pixels’ data in this manner, we generated a ‘correlation image’ from every subsequent stack of ten trials (7,990 frames), where each pixel’s value in the resulting image is the average correlation of the pixel’s fluorescence vector over 10 trials, with those of its 8 nearest neighbors. We then scaled the correlation image with the standard deviation of each pixels’ fluorescence over the ten-trial stack to enhance only those correlated pixels that also displayed large fluctuations in time. Following the generation of this sequence of statistical images, we used standard morphological operations (e.g. opening, dilation, erosion) to accentuate somatic shapes in these statistical images before seeding putative ROI centers as local maxima in these processed images. Pixels exhibiting high correlation within a 12 x 12 pixel bounding box around these local maxima then served as the ‘warm starts’ for a series of parallelized, local rank-1 nonnegative matrix factorizations centered on the patches of image data within these bounding boxes. Each image patch Y was thus factorized into the product of a ‘spatial footprint’ A and a temporal component C, with this matrix product added to the corresponding product of spatial and temporal background components B and f estimated over the same patch (*Y* = *AC* + *Bf*). These two background components are included to account for spatiotemporal sources of fluorescence variation present in the image patch that are nevertheless not neuronal in origin (e.g. out of focus fluorescence from neuropil activation in deeper/more superficial layers). Rank-1 solutions for A, C, B, and f are solvable with joint-convexity using the hierarchical alternating least squares algorithm. The resulting components A and C correspond to the neuronal shape (‘spatial filter’) and the demixed fluorescence timecourse of a neuron detected at the corresponding location. Following cell identification within the stacks, we used merging operations based on spatial- and temporal-overlap to merge redundantly-detected neurons in different 10-trial stacks across the full image dataset, to generate the final across-session matrix of cell identities and temporal footprints for all imaged trials.

#### Deconvolution of calcium transients

We deconvolved each neuron’s temporal component using the Online Active Set method to Infer Spikes (OASIS) algorithm (*45*), which simultaneously estimates the calcium signal of each neuron and the underlying spike train that gives rise to observed increases in calcium fluorescence. We used an iterative ‘spike-denoising’ procedure that leverages local fluorescence gradients centered around each inferred spike, as well as global statistics of each neuron’s fluorescence trace, to further exclude spikes detected spuriously by OASIS. To reintroduce the temporal autocorrelations characteristic of neuronal spike trains and reflect our uncertainty in the temporal precision of the spike-inference routine, we further re-convolved the denoised spike trains with a Gaussian kernel of standard deviation 143 ms (1.5 frames) and support 286 ms (3 frames). Unless otherwise specified, we used these smoothed spike trains for all further analyses.

### Calcium imaging data analyses

#### Identification of ‘active’ neurons by session

For each rat, spikes were estimated from calcium transients as described above, and used to calculate average firing rates of individual neurons, for all neurons detected across all 7 sessions. We computed separate average firing rates within the following periods of interest – (1) trial period: start to end of behavioral trial, (2) pre-lever period: trial start to lever presentation, (3) lever period: lever presentation to retraction, and (4) post-lever period: lever retraction to trial end. For each session, *a* neuron was deemed *active* in a period if its firing rate within that period was significantly different from that of the baseline period (5” window prior to start of the trial) using two-sided paired t-test. We corrected for multiple comparisons using Benjamini and Hochberg’s False Discovery Rate (FDR) method and set FDR at 0.05.

#### Quantification of shared vs. unique active neuron populations

For each rat, we identified *active* neurons in each period for all 7 imaging sessions, and used custom scripts to identify shared and unique *active* neuron populations between pairs of sessions. We used two measures to quantify the proportion of shared/unique *active* neuron populations between pairs of sessions (session A, session B):

1. %Overlap = number of neurons that were active in *both* Sessions A and B / number of neurons that were active in Sessions A or B = |A∩B| / |A∪B|
2. %Unique_A_ = number of neurons that were active *only in* Session A *but not* Session B / number of neurons that were active in Session A = |A-B| / |A|

For each rat, we also combined identified *active* neuron populations from the 3 session types (training, extinction, or reinstatement) and performed the same analysis, to identify the proportion of shared and unique *active* neuron populations between pairs of session-types.

#### Decoding trial-wise behavioral response from neuronal activity

For each rat, we trained linear decoders predicting the trial-wise behavioral outcome (response/no-response) from the vector of average firing rates of a set of neurons during a period of interest. We trained separate decoders for two distinct periods of interest: (1) lever period, and (2) pre-lever period. For each rat, we trained each decoder on a randomly selected subset of the data containing 75% of training and extinction trials. We reported the metrics resulting from applying the decoder to two separate test sets: (1) the remaining held-out 25% subset of training and extinction trials not used in training, and (2) all reinstatement trials unless otherwise stated. Note that no reinstatement trials were used in training the decoder. We resampled balanced samples (equal number of response and no-response trials) for the test set and reinstatement session in order to obtain a 50% chance level.

#### Contribution of Training and Extinction session active sub-populations to lever-period decoder accuracy

For each rat, we measured the lever period decoder accuracy in the absence of six specific *active* neuron sub-populations, to determine how much they contribute to decoder performance. The six sub-populations of *active* neurons considered were:

1. Training (T) – *active* in the 2 training sessions
2. Extinction (E) - *active* in the 4 extinction sessions
3. Training exclusive (T-E) – *active* in training but not extinction sessions
4. Extinction exclusive (E-T) – *active* in extinction but not training sessions
5. T or E (T ∪ E) – *active* in training, or extinction sessions
6. T & E (T ∩ E) – *active* in training and extinction sessions

We made the sub-population neurons absent in the decoder (already trained as described in the previous section) by setting their corresponding weights to 0. For each of the sub-populations, we also obtained a control accuracy by computing the average accuracy obtained leaving out randomly chosen sets of neurons of the same size. We used 2-way repeated measures ANOVA with within-subjects factors of *Population* (1-6) and *Drop type* (specific, random) to assess the effect of *active* population exclusion on overall decoder prediction accuracy. We used 2-way repeated measures ANOVA with within-subjects factors of *Drop type* (specific, random) and *Trial type* (response, no-response) to assess the effect of specific population exclusion on decoder prediction accuracy, and followed up with Bonferroni posthoc tests where appropriate (alpha level was set at 0.05).

## Author contributions

RM, DC, YS, and BTH conceptualized the project. RM, BL, GB, HJ, and DTL designed and constructed the experimental setup and imaging system. RM, DC, BLW, MV, LAR, CH, JMB, LEK, and SJW optimized surgical and imaging protocols, and performed behavioral and imaging experiments. RM, YZ, CH, JJ, KCL, KES, DJT, FP, and BTH developed analysis approaches. RM, YZ, CH, JJ, KCL, DJT, and KES performed data analysis. RM, CH, YZ, DJT, KES, YG, and DQP generated figures and statistical outputs. DC, BLW, MV, JMB, LAR, HJ, YS, GS, DTL, FP, and BTH contributed critical intellectual input throughout the project. RM, YZ, FP, and BTH wrote the manuscript with feedback from all authors.

## Acknowledgements

We thank the Genetically-Encoded Neuronal Indicator and Effector (GENIE) Project and the Janelia Research Campus of the Howard Hughes Medical Institute (HHMI) for generously allowing the use of GCaMP6 in our research. We thank members of the Hope and Shaham Lab (NIDA), and the Machine Learning Core (NIMH) for their feedback on study design and analysis, and for thoughtful comments during the writing of this manuscript. Research was supported by the NIDA Intramural Research Program (projects ZIA-DA000467, and ZIA-DA000434), and the NIMH Intramural Research Program (project ZIC-MH002968).

## Data, Materials, and Code Availability

All data generated or analyzed during this study, and needed to evaluate the conclusions in the paper, are included in the manuscript and supplementary materials. Raw imaging data and custom analysis scripts will be made available by the corresponding author upon reasonable request.

## Open Access

This article is licensed under a Creative Commons Attribution 4.0 International License, which permits use, sharing, adaptation, distribution, and reproduction in any medium or format, as long as you give appropriate credit to the original author(s) and the source, provide a link to the Creative Commons license, and indicate if changes were made. The images or other third-party material in this article are included in the article’s Creative Commons license, unless indicated otherwise in a credit line to the material. If material is not included in the article’s Creative Commons license and your intended use is not permitted by statutory regulation or exceeds the permitted use, you will need to obtain permission directly from the copyright holder. To view a copy of this license, visit http://creativecommons.org/licenses/by/4.0/.

© The Author(s) 2025

## Supplementary Figures

**Figure S1.**
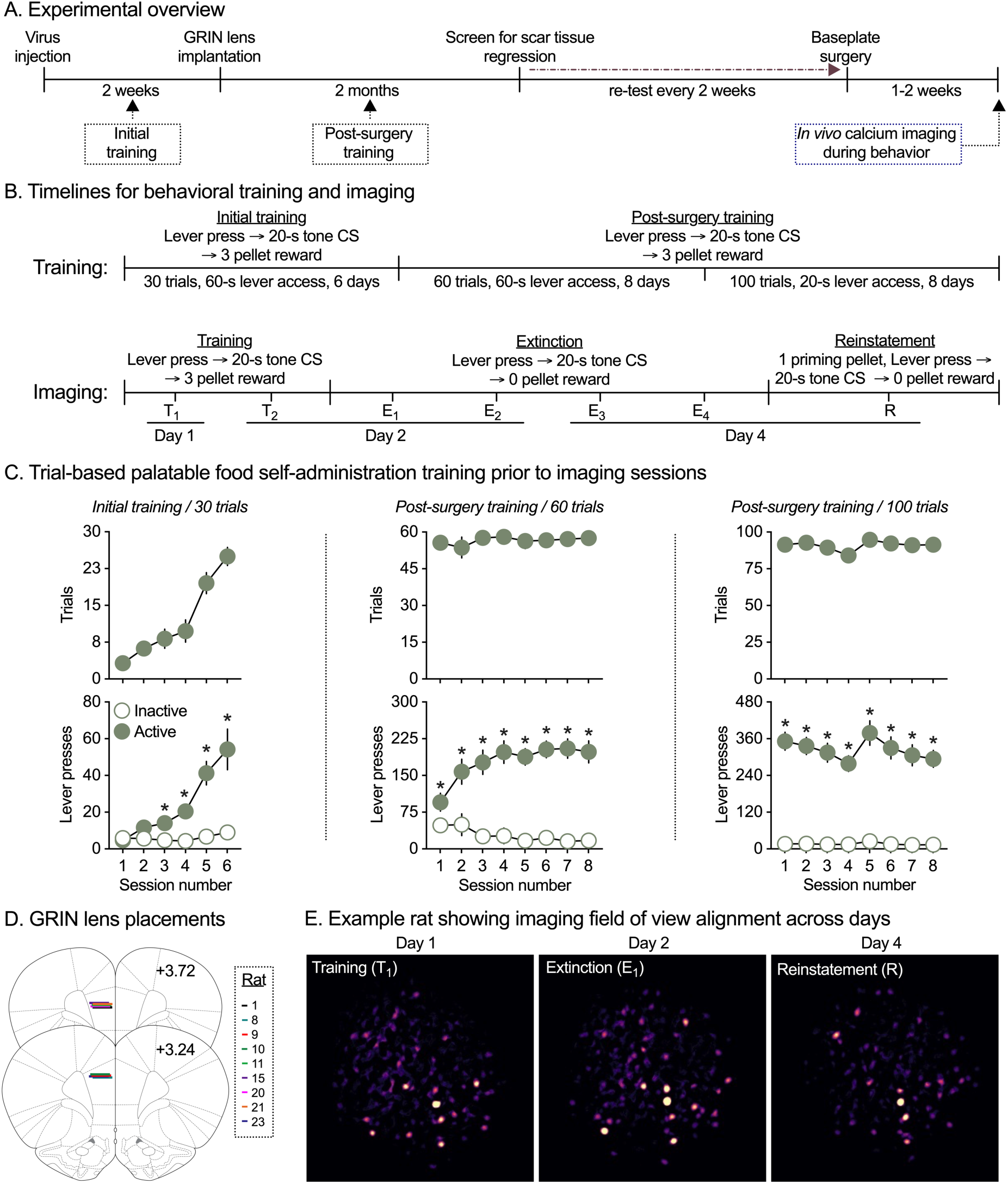
*In vivo* calcium imaging in rats during trial-based palatable food-seeking behavior. **(A)** *Experimental overview*. **(B)** *Timeline showing different surgical, behavioral training, and imaging stages*. **(C)** *Stages of trial-based palatable food self-administration training.* Rats were trained to self-administer palatable food pellets in three stages consisting of increasing numbers of trials and progressively shorter lever availability periods. Mean (± SEM) number of response trials (top row), and active and inactive lever presses (bottom row) during each training session for initial training (left panel, 30 trials with 60-s lever access/trial), post-surgery training phase 1 (center panel, 60 trials with 60-s lever access/trial) and post-surgery training phase 2 (right panel, 100 trials with 20-s lever access/trial). *Significant difference (*p* < 0.05) between active and inactive lever presses during a training session. **(D)** *GRIN lens placements.* Lines represent location of the base of imaging GRIN lens for rats included in the study (*n* = 9). **(E)** *Representative imaging field of view aligned across imaging days.* Maximum intensity projection of trial-wise correlation x standard deviation from one example rat to show alignment across sessions and over four imaging days.

**Figure S2.**
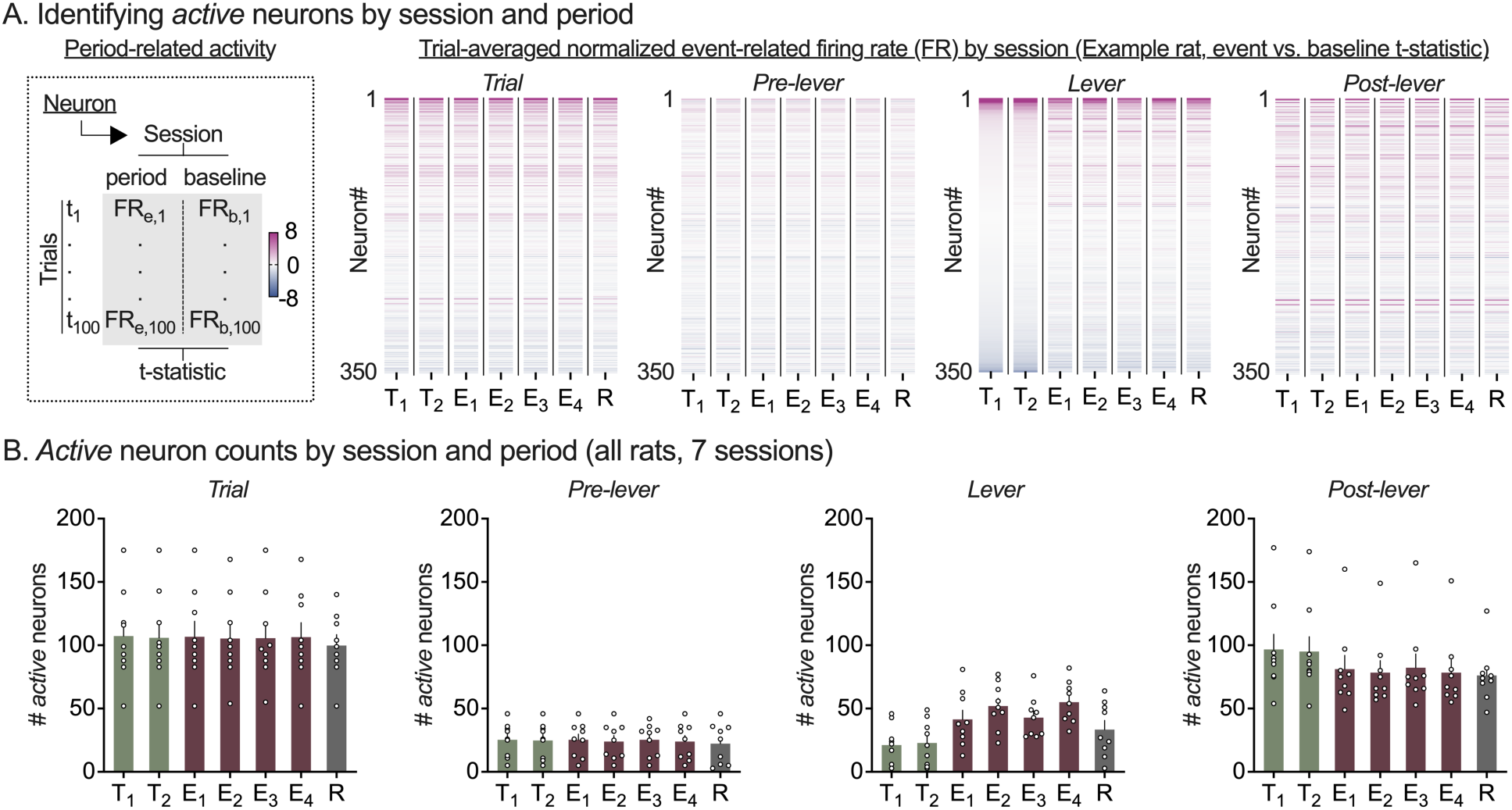
PL active neuron identification and overlap across all 7 imaging sessions. **(A)** *Active neuron identification within a trial period.* For each rat, spikes were estimated from calcium transients and used to calculate trial-by-trial average firing rate of individual neurons (all neurons detected across all 7 sessions) within each trial period. Next, for each session and trial period, *active* neurons were identified as those with significantly different average firing rate during that period vs. period at the start of the same trial in the session. Heatmaps showing t-statistic (period vs. baseline) of all neurons from one example rat for the entire trial (left), pre-lever period (center left), lever availability period (center right) and post-lever period (right). For all heatmaps, columns represent imaging session, and rows represent individual neurons ranked by t-statistic value for lever period activity in training session 1. **(B)** *Active neuron counts by trial period and imaging session.* Mean (± SEM) number of *active* neurons identified during the entire trial (left), pre-lever period (center left), lever availability period (center right) and post-lever period (right), for each imaging session. Clear circles represent data from individual rats (*n* = 9).

**Figure S3.**
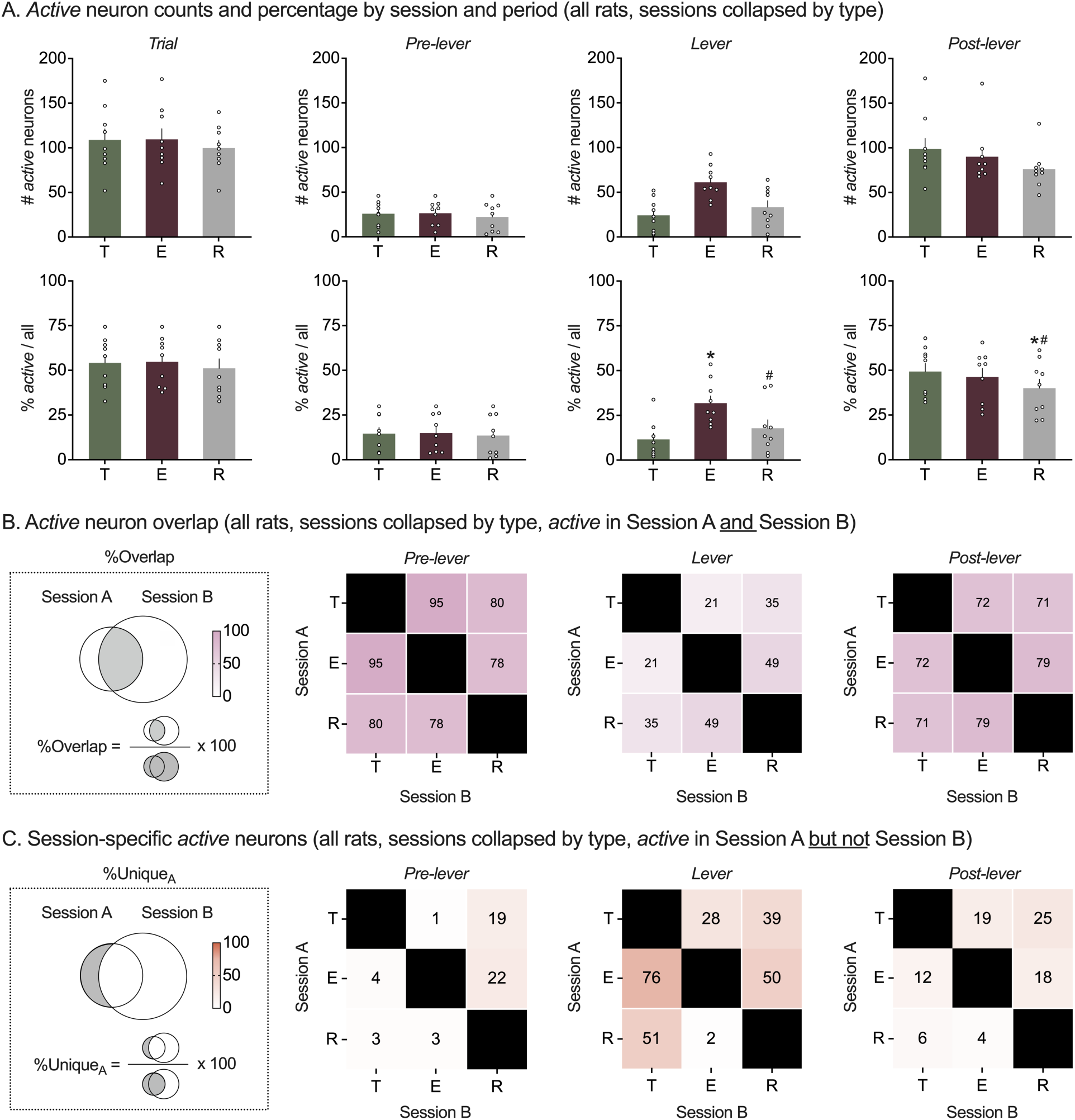
PL active neuron counts and overlap across 3 imaging session types (within trial period type). **(A)** *Active neuron identification trial period and imaging session. For each rat,* active neurons were detected within each trial period and for all 7 imaging sessions as described earlier and populations were then collapsed by 3 imaging session types (training, extinction or reinstatement). Mean (± SEM) number (top) and percentage (bottom) of *active* neurons identified during the entire trial (left), pre-lever period (center left), lever availability period (center right) and post-lever period (right), for each imaging session. Clear circles represent data from individual rats (*n* = 9). **(B)** Shared *active PL neurons across 3 imaging session types.* For each rat, within a trial period and session pair, the percentage of overlapping *active* neurons was calculated as shown in the schematic (left). Heatmaps show mean pairwise %overlapping neurons for pre-lever (left), lever period (middle) and post-lever period (right), for the 3 imaging session types. **(C)** S*ession-specific active PL neurons across 3 imaging session types.* For each rat, within a trial period and session type pair, the percentage of session-specific *active* neurons was calculated as shown in the schematic (left). Heatmaps show mean pairwise %unique neurons for pre-lever (left), lever period (middle) and post-lever period (right), for the 3 imaging session types.

**Figure S4.**
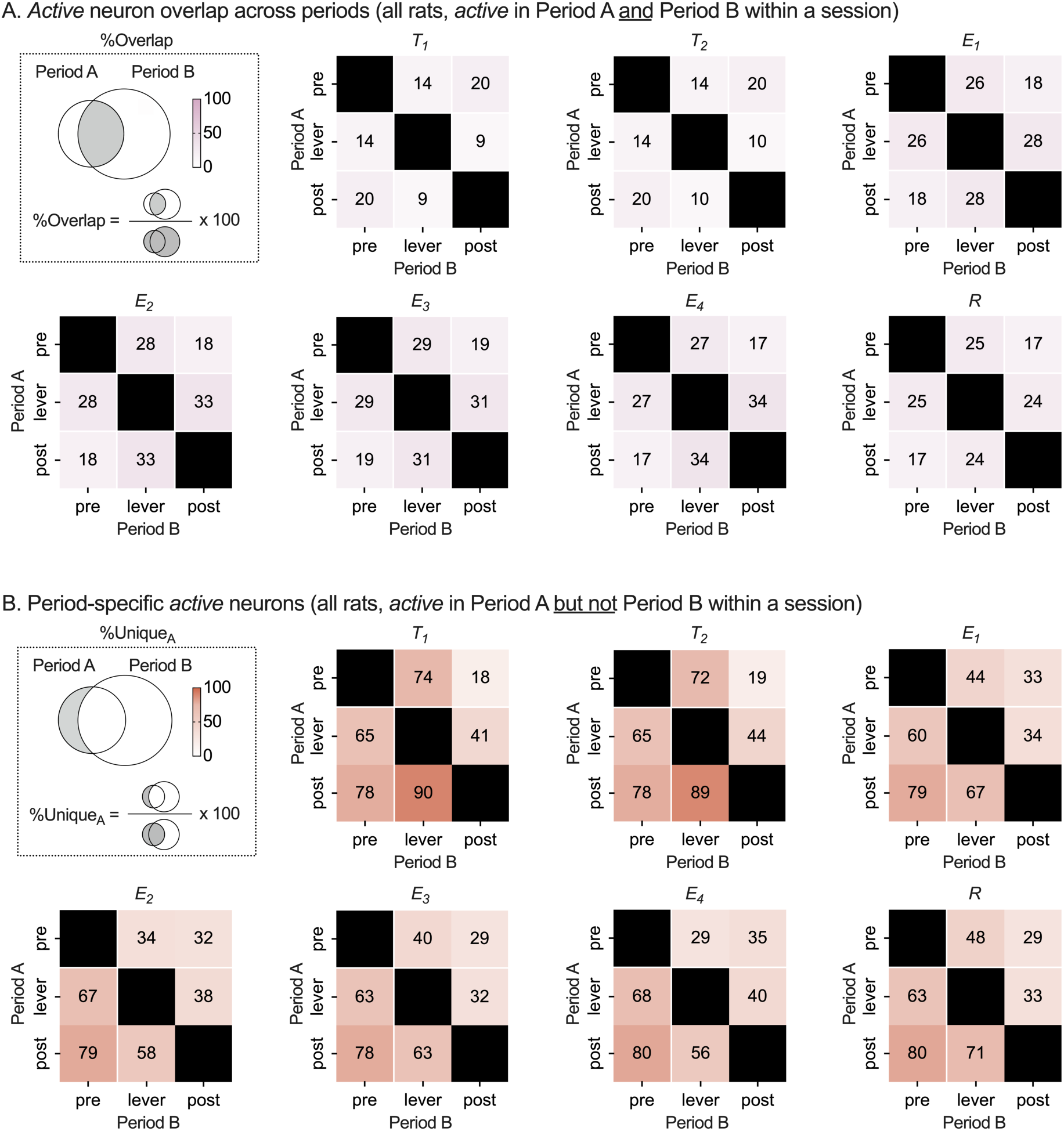
Shared and specific PL active neurons across trial periods (within imaging session). **(A)** *Active neuron overlap across trial periods. For each rat,* active neurons were detected within each trial period and for all 7 imaging sessions as described earlier. Next, within a session, percentage *active* neuron overlap was calculated pairwise between the three trial periods (pre-level, lever, post-lever) as shown in the schematic (left). Heatmaps show mean pairwise %overlap for each imaging session (7 sessions). **(B)** Period*-specific active PL neurons.* For each rat, within a trial period pair, the percentage of period-specific *active* neurons was calculated as shown in the schematic (left). Heatmaps show mean pairwise %unique neurons for each imaging session (7 sessions).

**Figure S5.**
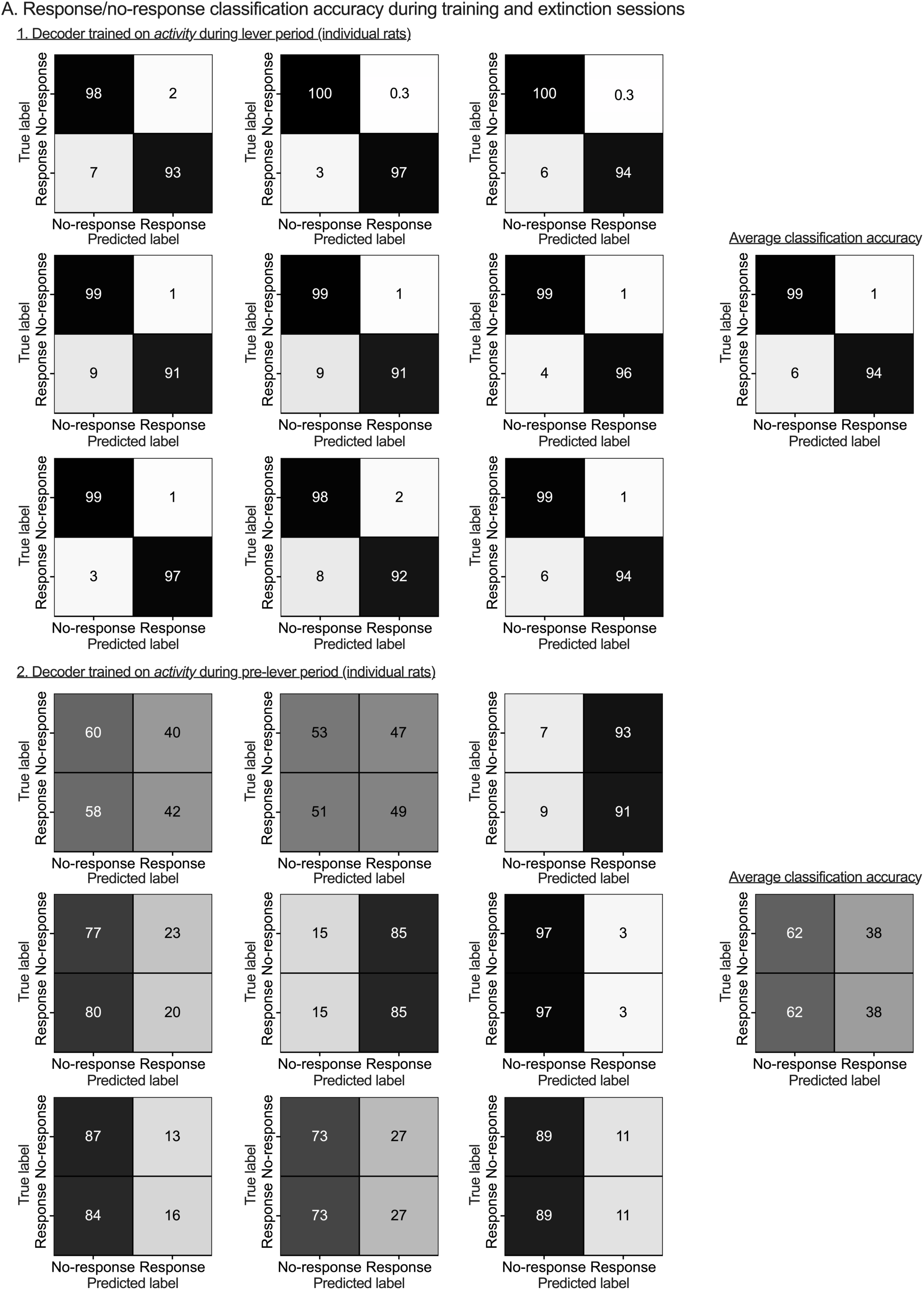
Individual rat lever period and pre-lever period decoder accuracy during training and extinction sessions. **(A.1)** Lever period decoder: For each rat, a binary (response/no-response) linear decoder was trained using the average firing rate vector during lever availability period, for a randomly selected subset of training and extinction trials (75/25 stratified split between training and test trials) and tested on the holdout subset of training and extinction trials. Response/no-response classification accuracy on the holdout test set (training and extinction) split by response and shown as a confusion matrix for individual rats (left 3 x 3 heatmap grid) or averaged across all rats (right heatmap, *n* = 9). **(A.2)** Pre-lever period decoder: For each rat, a binary (response/no-response) linear decoder was trained using the average firing rate vector during pre-lever availability period, for a randomly selected subset of training and extinction trials (75/25 stratified split between training and test trials) and tested on the holdout subset of training and extinction trials. Response/no-response classification accuracy on the holdout test set (training and extinction) split by response and shown as a confusion matrix for individual rats (left 3 x 3 heatmap grid) or averaged across all rats (right heatmap, *n* = 9).

**Figure S6.**
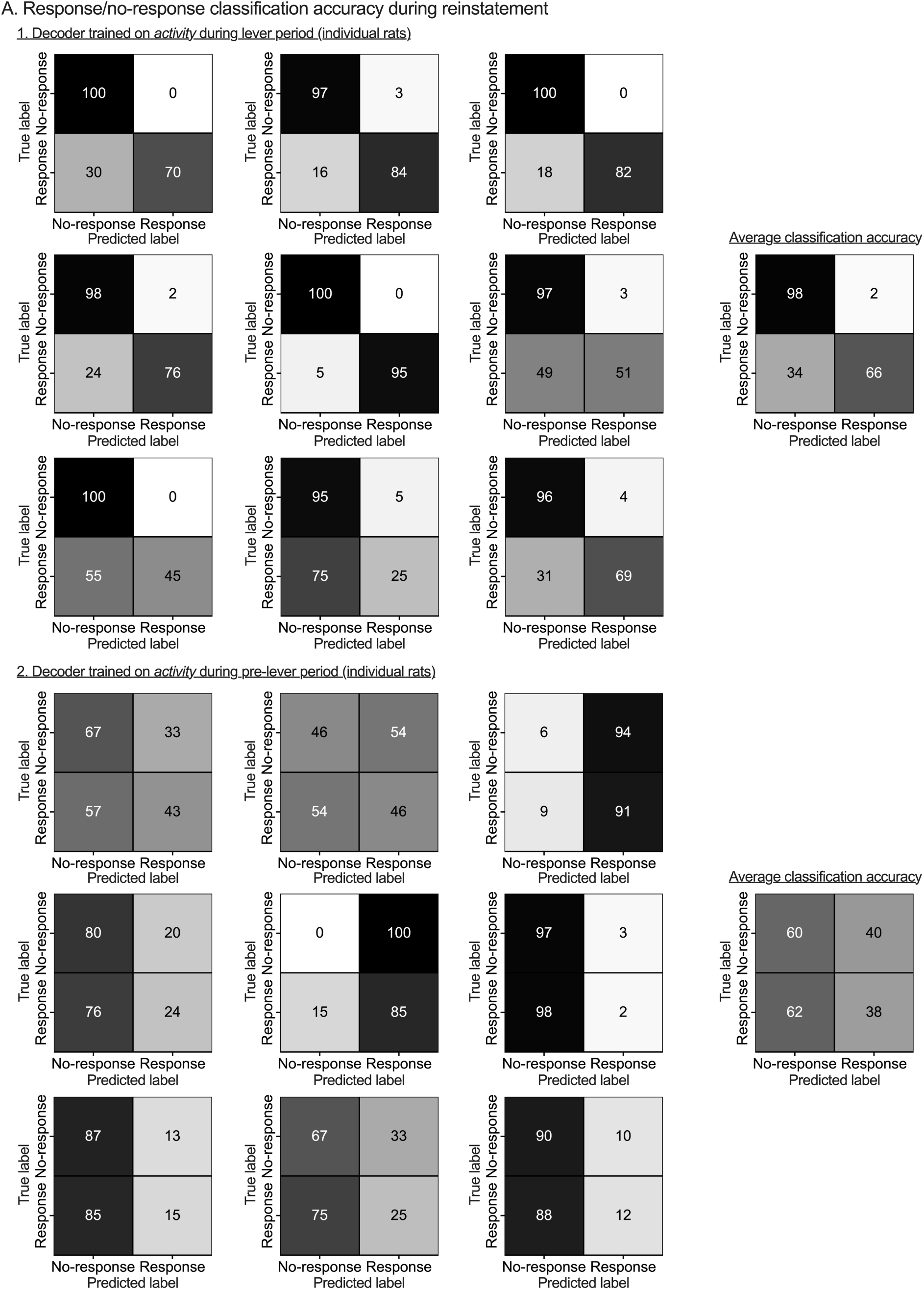
Individual rat lever period and pre-lever period decoder accuracy during the reinstatement session. **(A.1)** Lever period decoder: For each rat, a binary (response/no-response) linear decoder was trained using the average firing rate vector during lever availability period, for a randomly selected subset of training and extinction trials (75/25 stratified split between training and test trials) and tested on all reinstatement session trials. Response/no-response classification accuracy during the reinstatement session split by response and shown as a confusion matrix for individual rats (left 3 x 3 heatmap grid) or averaged across all rats (right heatmap, *n* = 9). **(A.2)** Pre-lever period decoder: For each rat, a binary (response/no-response) linear decoder was trained using the average firing rate vector during pre-lever availability period, for a randomly selected subset of training and extinction trials (75/25 stratified split between training and test trials) and tested on all reinstatement session trials. Response/no-response classification accuracy on the reinstatement session split by response and shown as a confusion matrix for individual rats (left 3 x 3 heatmap grid) or averaged across all rats (right heatmap, *n* = 9).

## Supplementary tables

**Table S1.**
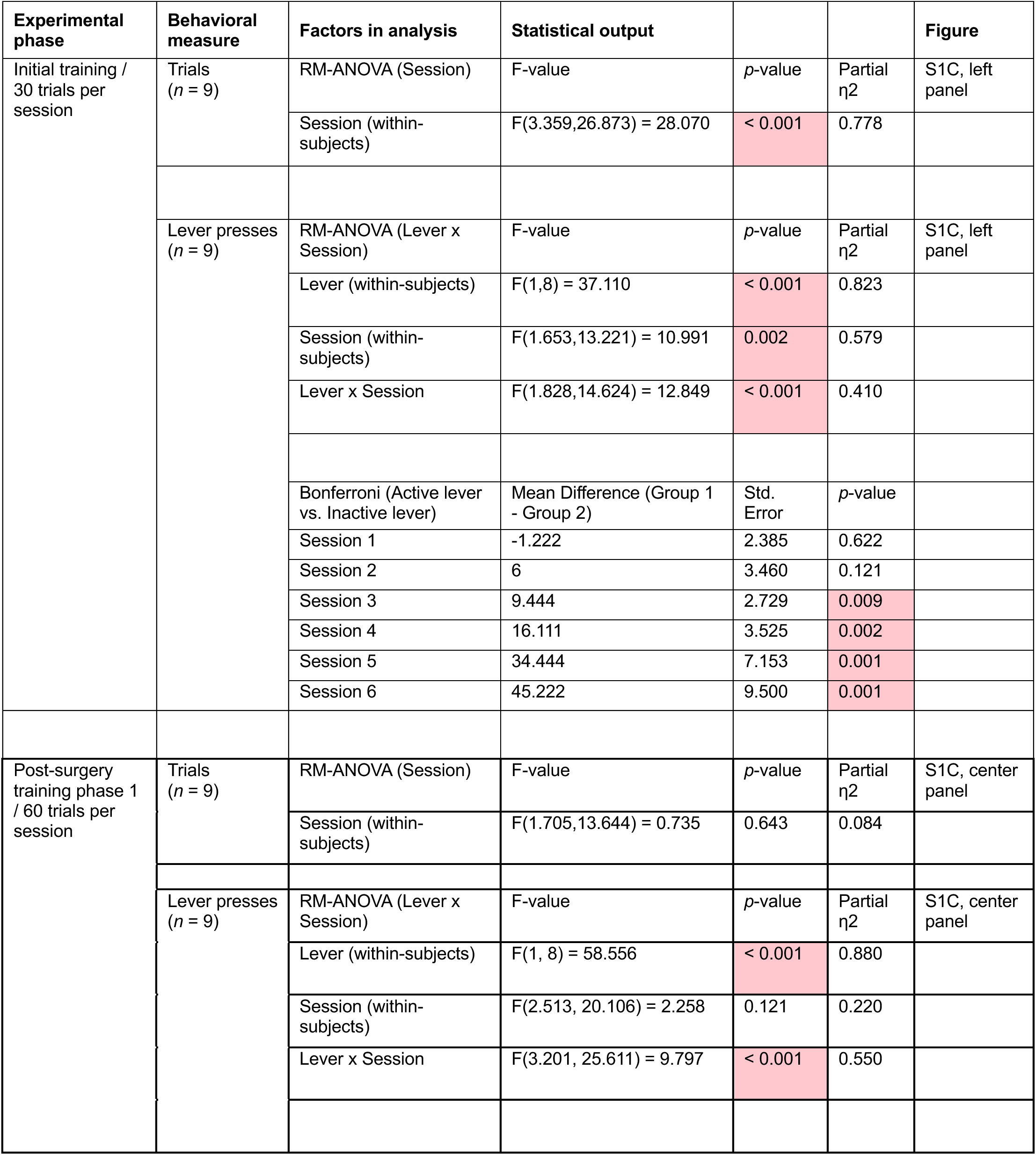
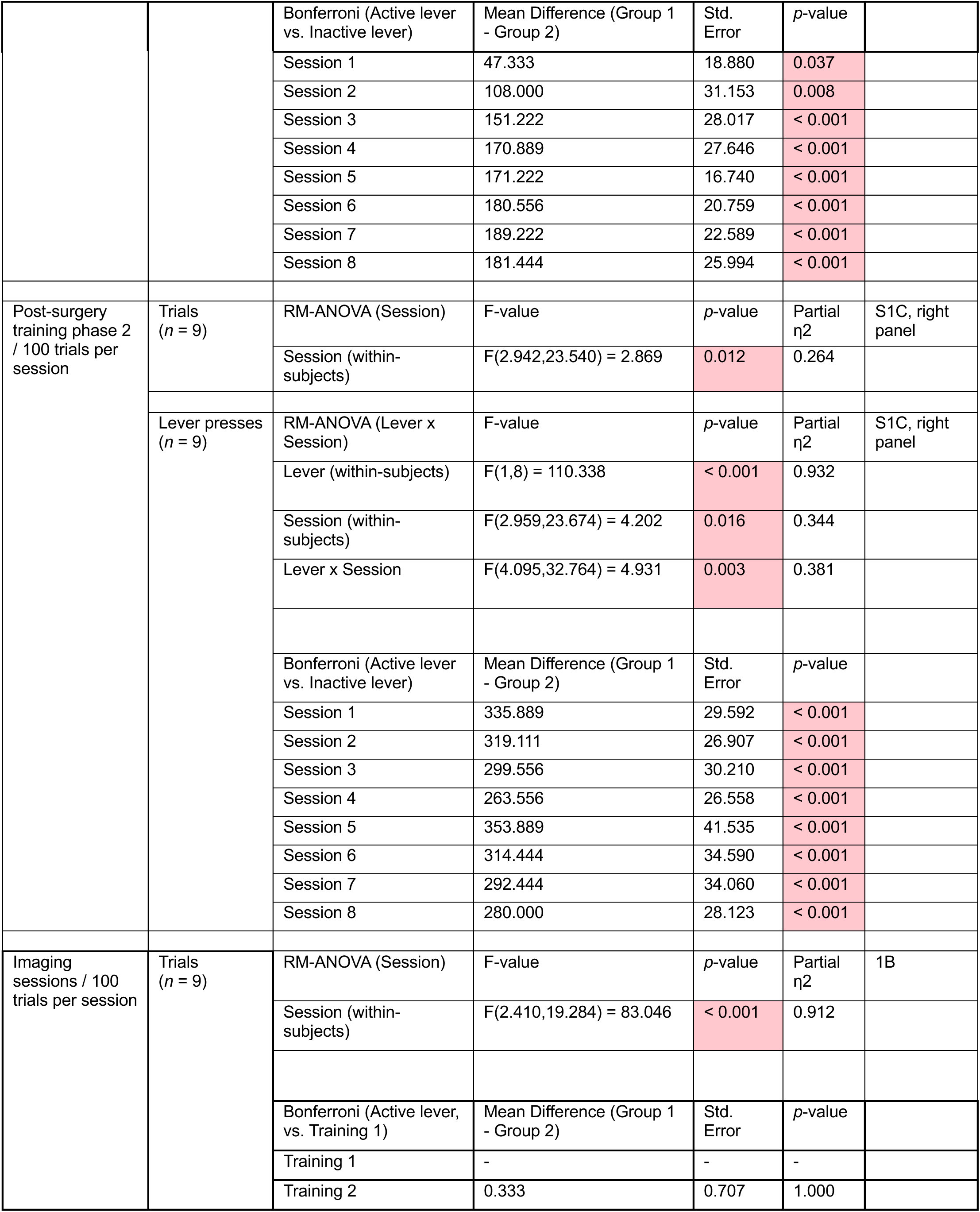
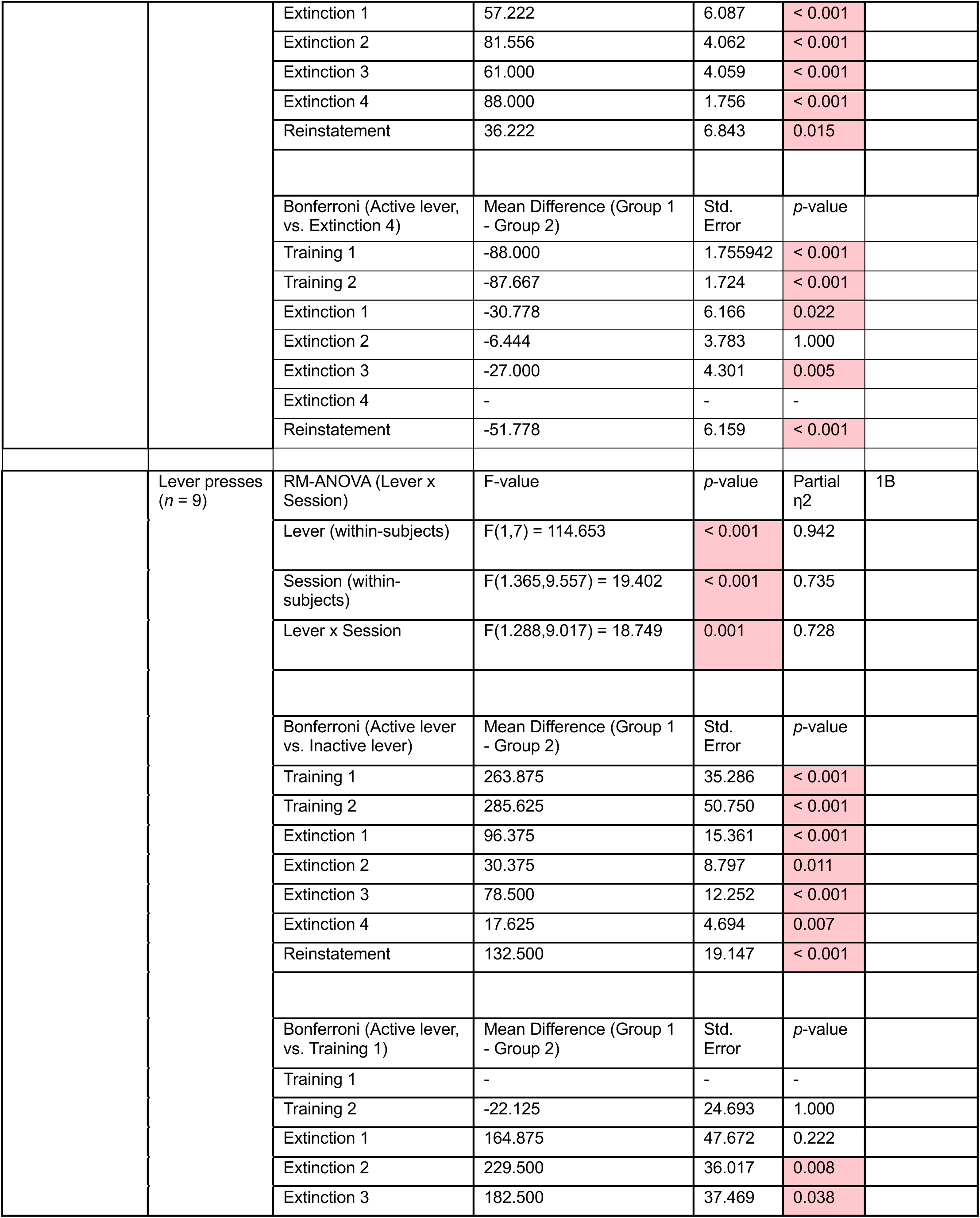
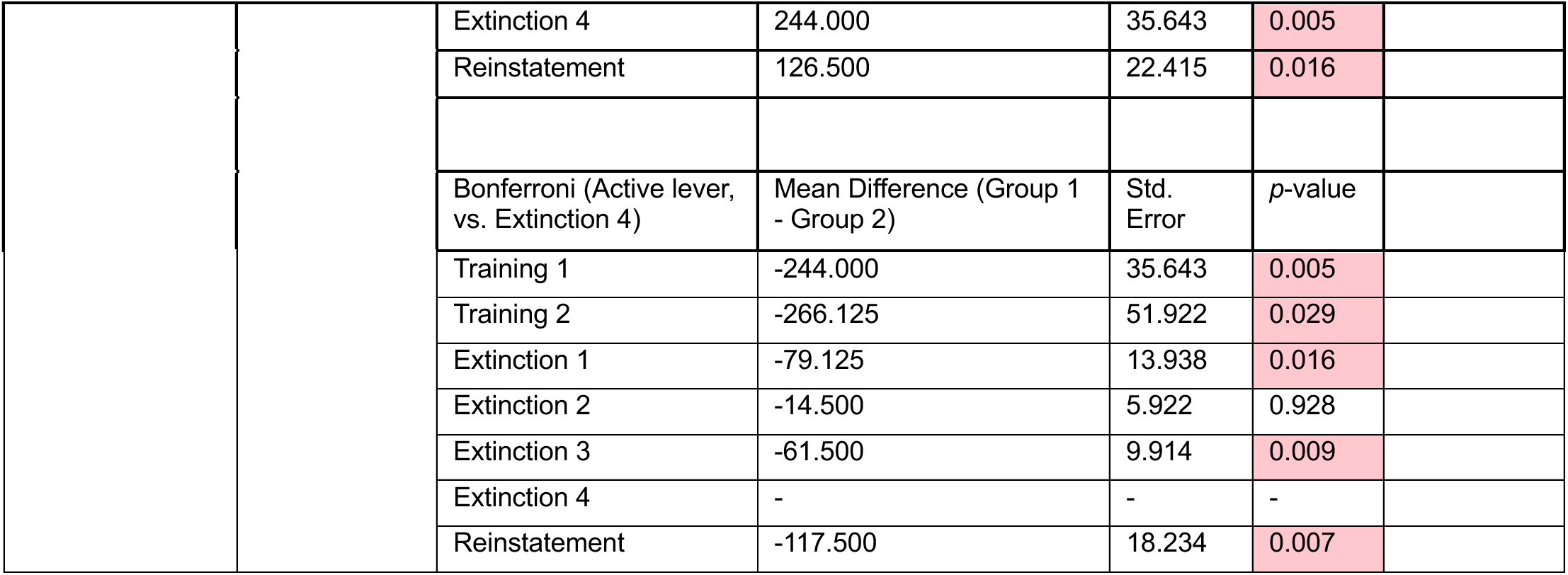
Detailed statistical outputs for behavior during training and imaging sessions.

| Experimental phase | Behavioral measure | Factors in analysis | Statistical output |  |  | Figure |
| --- | --- | --- | --- | --- | --- | --- |
| Initial training / 30 trials per session | Trials (n = 9) | RM-ANOVA (Session) | F-value | p-value | Partial $\eta^2$ | S1C, left panel |
|  |  | Session (within-subjects) | F(3.359,26.873) = 28.070 | < 0.001 | 0.778 |  |
| | Lever presses (n = 9) | RM-ANOVA (Lever x Session) | F-value | p-value | Partial $\eta^2$ | S1C, left panel |
|  |  | Lever (within-subjects) | F(1,8) = 37.110 | < 0.001 | 0.823 |  |
|  |  | Session (within-subjects) | F(1.653,13.221) = 10.991 | 0.002 | 0.579 |  |
|  |  | Lever x Session | F(1.828,14.624) = 12.849 | < 0.001 | 0.410 |  |
|  |  | Bonferroni (Active lever vs. Inactive lever) | Mean Difference (Group 1 - Group 2) | Std. Error | p-value |  |
|  |  | Session 1 | -1.222 | 2.385 | 0.622 |  |
|  |  | Session 2 | 6 | 3.460 | 0.121 |  |
|  |  | Session 3 | 9.444 | 2.729 | 0.009 |  |
|  |  | Session 4 | 16.111 | 3.525 | 0.002 |  |
|  |  | Session 5 | 34.444 | 7.153 | 0.001 |  |
|  |  | Session 6 | 45.222 | 9.500 | 0.001 |  |
| Post-surgery training phase 1 / 60 trials per session | Trials (n = 9) | RM-ANOVA (Session) | F-value | p-value | Partial $\eta^2$ | S1C, center panel |
|  |  | Session (within-subjects) | F(1.705,13.644) = 0.735 | 0.643 | 0.084 |  |
| | Lever presses (n = 9) | RM-ANOVA (Lever x Session) | F-value | p-value | Partial $\eta^2$ | S1C, center panel |
|  |  | Lever (within-subjects) | F(1, 8) = 58.556 | < 0.001 | 0.880 |  |
|  |  | Session (within-subjects) | F(2.513, 20.106) = 2.258 | 0.121 | 0.220 |  |
|  |  | Lever x Session | F(3.201, 25.611) = 9.797 | < 0.001 | 0.550 |  |
|  |  | Bonferroni (Active lever vs. Inactive lever) | Mean Difference (Group 1 - Group 2) | Std. Error | p-value |  |
|  |  | Session 1 | 47.333 | 18.880 | 0.037 |  |
|  |  | Session 2 | 108.000 | 31.153 | 0.008 |  |
|  |  | Session 3 | 151.222 | 28.017 | < 0.001 |  |
|  |  | Session 4 | 170.889 | 27.646 | < 0.001 |  |
|  |  | Session 5 | 171.222 | 16.740 | < 0.001 |  |
|  |  | Session 6 | 180.556 | 20.759 | < 0.001 |  |
|  |  | Session 7 | 189.222 | 22.589 | < 0.001 |  |
|  |  | Session 8 | 181.444 | 25.994 | < 0.001 |  |
| Post-surgery training phase 2 / 100 trials per session | Trials (n = 9) | RM-ANOVA (Session) | F-value | p-value | Partial $\eta^2$ | S1C, right panel |
|  |  | Session (within-subjects) | F(2.942,23.540) = 2.869 | 0.012 | 0.264 |  |
| | Lever presses (n = 9) | RM-ANOVA (Lever x Session) | F-value | p-value | Partial $\eta^2$ | S1C, right panel |
|  |  | Lever (within-subjects) | F(1,8) = 110.338 | < 0.001 | 0.932 |  |
|  |  | Session (within-subjects) | F(2.959,23.674) = 4.202 | 0.016 | 0.344 |  |
|  |  | Lever x Session | F(4.095,32.764) = 4.931 | 0.003 | 0.381 |  |
|  |  | Bonferroni (Active lever vs. Inactive lever) | Mean Difference (Group 1 - Group 2) | Std. Error | p-value |  |
|  |  | Session 1 | 335.889 | 29.592 | < 0.001 |  |
|  |  | Session 2 | 319.111 | 26.907 | < 0.001 |  |
|  |  | Session 3 | 299.556 | 30.210 | < 0.001 |  |
|  |  | Session 4 | 263.556 | 26.558 | < 0.001 |  |
|  |  | Session 5 | 353.889 | 41.535 | < 0.001 |  |
|  |  | Session 6 | 314.444 | 34.590 | < 0.001 |  |
|  |  | Session 7 | 292.444 | 34.060 | < 0.001 |  |
|  |  | Session 8 | 280.000 | 28.123 | < 0.001 |  |
| Imaging sessions / 100 trials per session | Trials (n = 9) | RM-ANOVA (Session) | F-value | p-value | Partial $\eta^2$ | 1B |
|  |  | Session (within-subjects) | F(2.410,19.284) = 83.046 | < 0.001 | 0.912 |  |
|  |  | Bonferroni (Active lever, vs. Training 1) | Mean Difference (Group 1 - Group 2) | Std. Error | p-value |  |
|  |  | Training 1 | - | - | - |  |
|  |  | Training 2 | 0.333 | 0.707 | 1.000 |  |
|  |  | Extinction 1 | 57.222 | 6.087 | < 0.001 |  |
|  |  | Extinction 2 | 81.556 | 4.062 | < 0.001 |  |
|  |  | Extinction 3 | 61.000 | 4.059 | < 0.001 |  |
|  |  | Extinction 4 | 88.000 | 1.756 | < 0.001 |  |
|  |  | Reinstatement | 36.222 | 6.843 | 0.015 |  |
|  |  | Bonferroni (Active lever, vs. Extinction 4) | Mean Difference (Group 1 - Group 2) | Std. Error | p-value |  |
|  |  | Training 1 | -88.000 | 1.755942 | < 0.001 |  |
|  |  | Training 2 | -87.667 | 1.724 | < 0.001 |  |
|  |  | Extinction 1 | -30.778 | 6.166 | 0.022 |  |
|  |  | Extinction 2 | -6.444 | 3.783 | 1.000 |  |
|  |  | Extinction 3 | -27.000 | 4.301 | 0.005 |  |
|  |  | Extinction 4 | - | - | - |  |
|  |  | Reinstatement | -51.778 | 6.159 | < 0.001 |  |
| | Lever presses<br>(n = 9) | RM-ANOVA (Lever x Session) | F-value | p-value | Partial $\eta^2$ | 1B |
|  |  | Lever (within-subjects) | F(1,7) = 114.653 | < 0.001 | 0.942 |  |
|  |  | Session (within-subjects) | F(1.365,9.557) = 19.402 | < 0.001 | 0.735 |  |
|  |  | Lever x Session | F(1.288,9.017) = 18.749 | 0.001 | 0.728 |  |
|  |  | Bonferroni (Active lever vs. Inactive lever) | Mean Difference (Group 1 - Group 2) | Std. Error | p-value |  |
|  |  | Training 1 | 263.875 | 35.286 | < 0.001 |  |
|  |  | Training 2 | 285.625 | 50.750 | < 0.001 |  |
|  |  | Extinction 1 | 96.375 | 15.361 | < 0.001 |  |
|  |  | Extinction 2 | 30.375 | 8.797 | 0.011 |  |
|  |  | Extinction 3 | 78.500 | 12.252 | < 0.001 |  |
|  |  | Extinction 4 | 17.625 | 4.694 | 0.007 |  |
|  |  | Reinstatement | 132.500 | 19.147 | < 0.001 |  |
|  |  | Bonferroni (Active lever, vs. Training 1) | Mean Difference (Group 1 - Group 2) | Std. Error | p-value |  |
|  |  | Training 1 | - | - | - |  |
|  |  | Training 2 | -22.125 | 24.693 | 1.000 |  |
|  |  | Extinction 1 | 164.875 | 47.672 | 0.222 |  |
|  |  | Extinction 2 | 229.500 | 36.017 | 0.008 |  |
|  |  | Extinction 3 | 182.500 | 37.469 | 0.038 |  |

|  |  |  |  |  |
| --- | --- | --- | --- | --- |
|  | Extinction 4 | 244.000 | 35.643 | 0.005 |
|  | Reinstatement | 126.500 | 22.415 | 0.016 |
|  | Bonferroni (Active lever, vs. Extinction 4) | Mean Difference (Group 1 - Group 2) | Std. Error | p-value |
|  | Training 1 | -244.000 | 35.643 | 0.005 |
|  | Training 2 | -266.125 | 51.922 | 0.029 |
|  | Extinction 1 | -79.125 | 13.938 | 0.016 |
|  | Extinction 2 | -14.500 | 5.922 | 0.928 |
|  | Extinction 3 | -61.500 | 9.914 | 0.009 |
|  | Extinction 4 | - | - | - |
|  | Reinstatement | -117.500 | 18.234 | 0.007 |

**Table S2.**
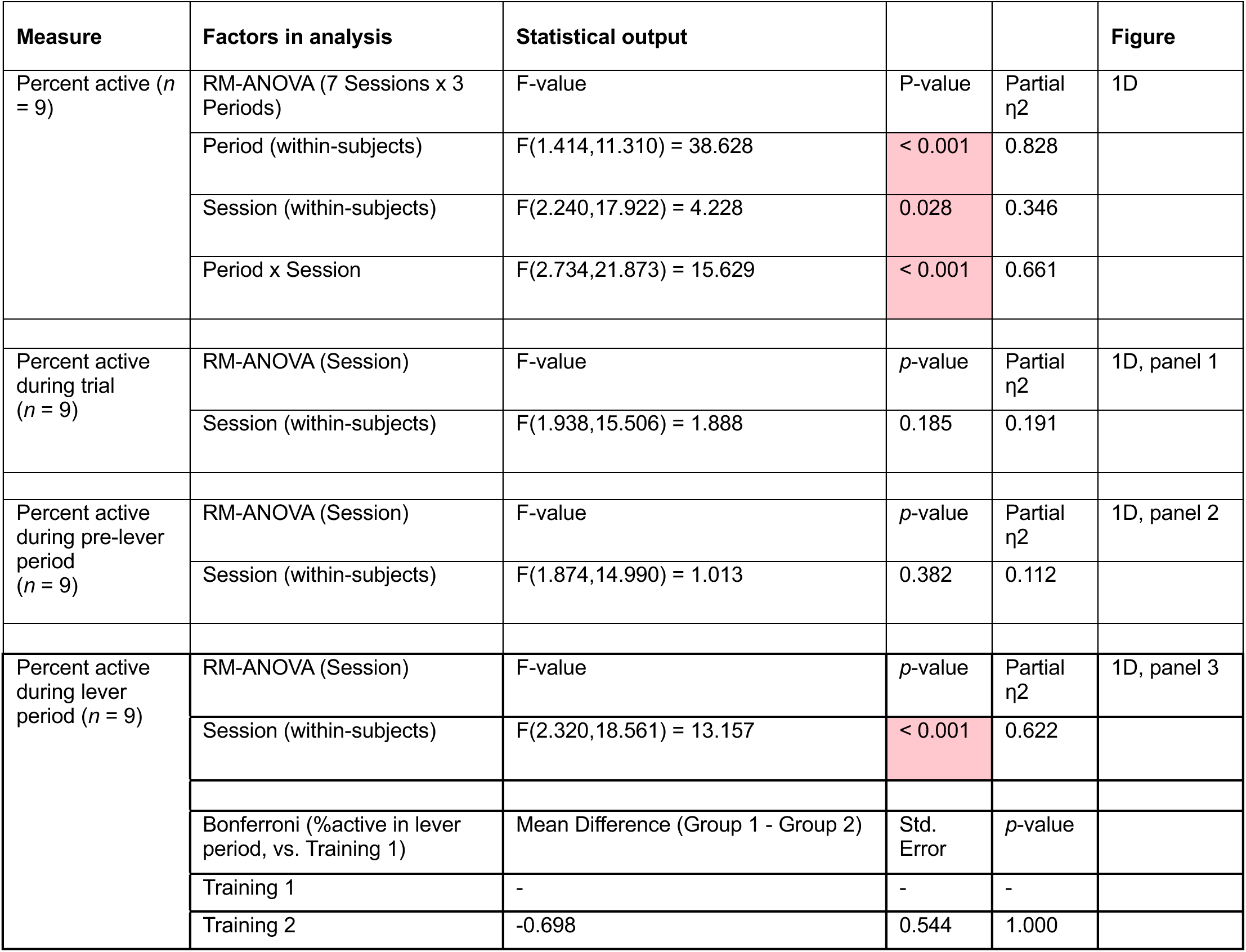
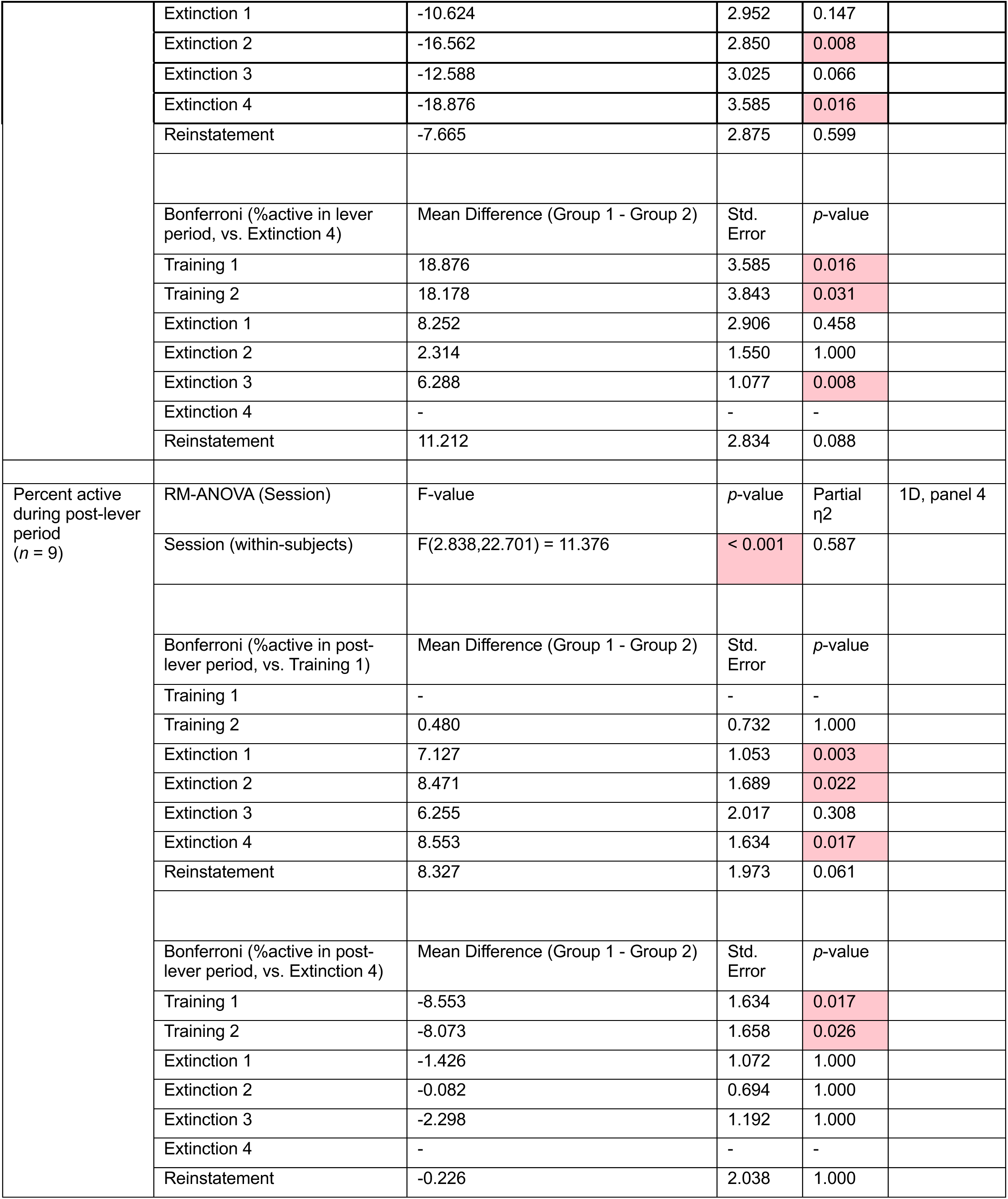
Detailed statistical outputs for analyses pertaining to number and percentage of active neurons by session (7 sessions) and trial period.

| Measure | Factors in analysis | Statistical output |  |  | Figure |
| --- | --- | --- | --- | --- | --- |
| Percent active ($n = 9$) | RM-ANOVA (7 Sessions x 3 Periods) | F-value | P-value | Partial $\eta^2$ | 1D |
| | Period (within-subjects) | $F(1.414, 11.310) = 38.628$ | < 0.001 | 0.828 | |
| | Session (within-subjects) | $F(2.240, 17.922) = 4.228$ | 0.028 | 0.346 | |
| | Period x Session | $F(2.734, 21.873) = 15.629$ | < 0.001 | 0.661 | |
| Percent active during trial ($n = 9$) | RM-ANOVA (Session) | F-value | p-value | Partial $\eta^2$ | 1D, panel 1 |
| | Session (within-subjects) | $F(1.938, 15.506) = 1.888$ | 0.185 | 0.191 | |
| Percent active during pre-lever period ($n = 9$) | RM-ANOVA (Session) | F-value | p-value | Partial $\eta^2$ | 1D, panel 2 |
| | Session (within-subjects) | $F(1.874, 14.990) = 1.013$ | 0.382 | 0.112 | |
| Percent active during lever period ($n = 9$) | RM-ANOVA (Session) | F-value | p-value | Partial $\eta^2$ | 1D, panel 3 |
| | Session (within-subjects) | $F(2.320, 18.561) = 13.157$ | < 0.001 | 0.622 | |
|  | Bonferroni (%active in lever period, vs. Training 1) | Mean Difference (Group 1 - Group 2) | Std. Error | p-value |  |
|  | Training 1 | - | - | - |  |
|  | Training 2 | -0.698 | 0.544 | 1.000 |  |
|  | Extinction 1 | -10.624 | 2.952 | 0.147 |  |
|  | Extinction 2 | -16.562 | 2.850 | 0.008 |  |
|  | Extinction 3 | -12.588 | 3.025 | 0.066 |  |
|  | Extinction 4 | -18.876 | 3.585 | 0.016 |  |
|  | Reinstatement | -7.665 | 2.875 | 0.599 |  |
|  | Bonferroni (%active in lever period, vs. Extinction 4) | Mean Difference (Group 1 - Group 2) | Std. Error | *p*-value |  |
|  | Training 1 | 18.876 | 3.585 | 0.016 |  |
|  | Training 2 | 18.178 | 3.843 | 0.031 |  |
|  | Extinction 1 | 8.252 | 2.906 | 0.458 |  |
|  | Extinction 2 | 2.314 | 1.550 | 1.000 |  |
|  | Extinction 3 | 6.288 | 1.077 | 0.008 |  |
|  | Extinction 4 | - | - | - |  |
|  | Reinstatement | 11.212 | 2.834 | 0.088 |  |
| Percent active during post-lever period (*n* = 9) | RM-ANOVA (Session) | F-value | *p*-value | Partial $\eta^2$ | 1D, panel 4 |
|  | Session (within-subjects) | F(2.838,22.701) = 11.376 | < 0.001 | 0.587 |  |
|  | Bonferroni (%active in post-lever period, vs. Training 1) | Mean Difference (Group 1 - Group 2) | Std. Error | *p*-value |  |
|  | Training 1 | - | - | - |  |
|  | Training 2 | 0.480 | 0.732 | 1.000 |  |
|  | Extinction 1 | 7.127 | 1.053 | 0.003 |  |
|  | Extinction 2 | 8.471 | 1.689 | 0.022 |  |
|  | Extinction 3 | 6.255 | 2.017 | 0.308 |  |
|  | Extinction 4 | 8.553 | 1.634 | 0.017 |  |
|  | Reinstatement | 8.327 | 1.973 | 0.061 |  |
|  | Bonferroni (%active in post-lever period, vs. Extinction 4) | Mean Difference (Group 1 - Group 2) | Std. Error | *p*-value |  |
|  | Training 1 | -8.553 | 1.634 | 0.017 |  |
|  | Training 2 | -8.073 | 1.658 | 0.026 |  |
|  | Extinction 1 | -1.426 | 1.072 | 1.000 |  |
|  | Extinction 2 | -0.082 | 0.694 | 1.000 |  |
|  | Extinction 3 | -2.298 | 1.192 | 1.000 |  |
|  | Extinction 4 | - | - | - |  |
|  | Reinstatement | -0.226 | 2.038 | 1.000 |  |

**Table S3.** Detailed statistical outputs for analyses pertaining to number and percentage of active neurons by session-type (3 session types) and trial period.

| Measure | Factors in analysis | Statistical output |  |  | Figure |
| --- | --- | --- | --- | --- | --- |
| Percent active during trial ($n = 9$) | RM-ANOVA (3 Session types x 3 Periods) | F-value | $p$-value | Partial $\eta^2$ | S3A |
| | Period (within-subjects) | $F(1.494, 11.956) = 37.606$ | < 0.001 | 0.825 | |
| | Session type (within-subjects) | $F(1.304, 10.429) = 18.018$ | < 0.001 | 0.693 | |
| | Period x Session type | $F(2.325, 18.599) = 18.933$ | < 0.001 | 0.703 | |
| Percent active during trial ($n = 9$) | RM-ANOVA (Session type) | F-value | $p$-value | Partial $\eta^2$ | S3A, panel 1 |
| | Session (within-subjects) | $F(1.889, 15.111) = 3.888$ | 0.042 | 0.327 | |
| | Bonferroni (%active in trial, vs. Training) | Mean Difference (Group 1 - Group 2) | Std. Error | $p$-value | |
|  | Training | - | - | - |  |
|  | Extinction | -0.580 | 1.243 | 1.000 |  |
|  | Reinstatement | 3.000 | 1.527 | 0.255 |  |
| | Bonferroni (%active in trial, vs. Extinction) | Mean Difference (Group 1 - Group 2) | Std. Error | $p$-value | |
|  | Training | 0.580 | 1.243 | 1.000 |  |
|  | Extinction | - | - | - |  |
|  | Reinstatement | 3.580 | 1.350 | 0.087 |  |
| Percent active during pre-lever period ($n = 9$) | RM-ANOVA (Session type) | F-value | $p$-value | Partial $\eta^2$ | |
| | Session (within-subjects) | $F(1.437, 11.499) = 3.139$ | 0.071 | 0.282 | S3A, panel 2 |
| Percent active during lever period ($n = 9$) | RM-ANOVA (Session type) | F-value | $p$-value | Partial $\eta^2$ | S3A, panel 3 |
| | Session (within-subjects) | $F(1.679, 13.433) = 22.626$ | < 0.001 | 0.739 | |
| | Bonferroni (%active in lever period, vs. Training) | Mean Difference (Group 1 - Group 2) | Std. Error | $p$-value | |
|  | Training | - | - | - |  |
|  | Extinction | -20.194 | 3.635 | 0.002 |  |
|  | Reinstatement | -6.173 | 3.018 | 0.225 |  |
| | Bonferroni (%active in lever period, vs. Extinction) | Mean Difference (Group 1 - Group 2) | Std. Error | $p$-value | |
|  | Training | -20.194 | 3.635 | 0.002 |  |
|  | Extinction | - | - | - |  |
|  | Reinstatement | 14.020 | 2.465 | 0.001 |  |
| Percent active during post-lever period ($n = 9$) | RM-ANOVA (Session type) | F-value | $p$-value | Partial $\eta^2$ | S3A, panel 4 |
| | Session (within-subjects) | $F(1.947, 15.574) = 10.909$ | 0.001 | 0.577 | |
| | Bonferroni (%active in post-lever period, vs. Training) | Mean Difference (Group 1 - Group 2) | Std. Error | $p$-value | |
|  | Training | - | - | - |  |
|  | Extinction | 3.077 | 2.170 | 0.582 |  |
|  | Reinstatement | 9.314 | 1.877 | 0.003 |  |
| | Bonferroni (%active in post-lever period, vs. Extinction) | Mean Difference (Group 1 - Group 2) | Std. Error | $p$-value | |
|  | Training | 3.077 | 2.170 | 0.582 |  |
|  | Extinction | - | - | - |  |
|  | Reinstatement | 6.238 | 2.038 | 0.047 |  |

**Table S4.**
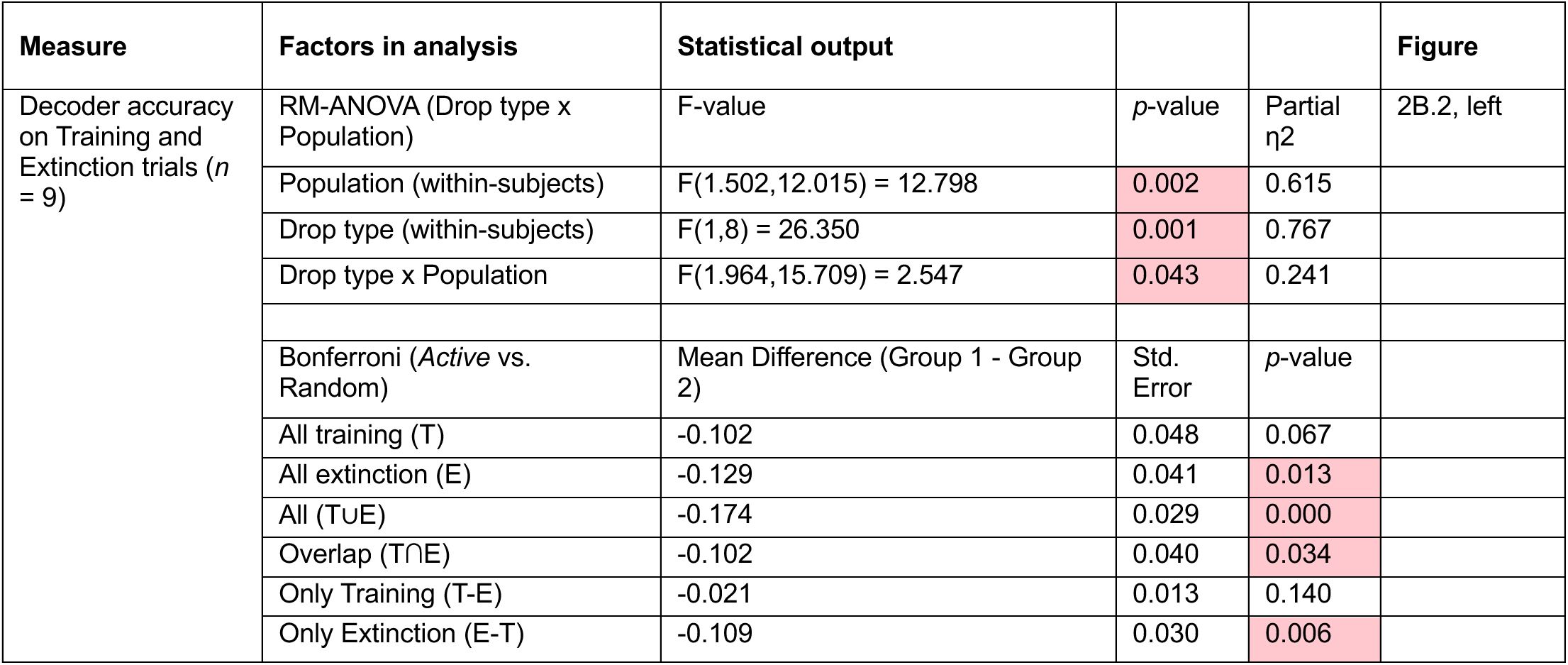
Detailed statistical outputs for trial-wise decoder accuracy during training and extinction sessions.

**Table S5.** Detailed statistical outputs for trial-wise decoder accuracy during training and extinction following exclusion of specific active neuron sub-populations.

| Measure | Active neuron population | Factors in analysis | Statistical output |  |  | Figure |
| --- | --- | --- | --- | --- | --- | --- |
| Decoder accuracy on Training and Extinction trials ($n = 9$) | All training (T) | RM-ANOVA (Response x Drop type) | F-value | $p$-value | Partial $\eta^2$ | 2B.3, top left |
| | | Response (within-subjects) | $F(1,8) = 6.209$ | 0.037 | 0.437 | |
| | | Drop type (within-subjects) | $F(1,8) = 3.977$ | 0.081 | 0.332 | |
| | | Response x Drop type | $F(1,8) = 6.088$ | 0.039 | 0.432 | |
| | | Bonferroni (*Active* vs. Random) | Mean Difference (Group 1 - Group 2) | Std. Error | $p$-value | |
|  |  | Response | -0.243 | 0.108 | 0.055 |  |
|  |  | No-response | 0.028 | 0.010 | 0.020 |  |
| | All extinction (E) | RM-ANOVA (Response x Drop type) | F-value | $p$-value | Partial $\eta^2$ | 2B.3, bottom left |
| | | Response (within-subjects) | $F(1,8) = 0.434$ | 0.529 | 0.051 | |
| | | Drop type (within-subjects) | $F(1,8) = 10.403$ | 0.012 | 0.565 | |
| | | Response x Drop type | $F(1,8) = 1.126$ | 0.320 | 0.123 | |
| | | Bonferroni (*Active* vs. Random) | Mean Difference (Group 1 - Group 2) | Std. Error | $p$-value | |
|  |  | Response | -0.059 | 0.092 | 0.540 |  |
|  |  | No-response | -0.246 | 0.108 | 0.051 |  |
| | Only Training (T-E) | RM-ANOVA (Response x Drop type) | F-value | $p$-value | Partial $\eta^2$ | 2B.3, top center |
| | | Response (within-subjects) | $F(1,8) = 0.278$ | 0.612 | 0.034 | |
| | | Drop type (within-subjects) | $F(1,8) = 2.993$ | 0.122 | 0.272 | |
| | | Response x Drop type | $F(1,8) = 1.901$ | 0.205 | 0.192 | |
| | | Bonferroni (*Active* vs. Random) | Mean Difference (Group 1 - Group 2) | Std. Error | $p$-value | |
|  |  | Response | 0.011 | 0.015 | 0.508 |  |
|  |  | No-response | -0.053 | 0.033 | 0.153 |  |
| | Only Extinction (E-T) | RM-ANOVA (Response x Drop type) | F-value | $p$-value | Partial $\eta^2$ | 2B.3, bottom center |
| | | Response (within-subjects) | $F(1,8) = 9.909$ | 0.014 | 0.553 | |
| | | Drop type (within-subjects) | $F(1,8) = 11.554$ | 0.009 | 0.591 | |
| | | Response x Drop type | $F(1,8) = 45.604$ | < 0.001 | 0.851 | |
| | | Bonferroni (*Active* vs. Random) | Mean Difference (Group 1 - Group 2) | Std. Error | $p$-value | |
|  |  | Response | 0.124 | 0.006 | <0.001 |  |
|  |  | No-response | -0.374 | 0.073 | <0.001 |  |
| | Either T or E (T∪E) | RM-ANOVA (Response x Drop type) | F-value | p-value | Partial $\eta^2$ | 2B.3, top right |
|  |  | Response (within-subjects) | F(1,8) = 0.450 | 0.521 | 0.053 |  |
|  |  | Drop type (within-subjects) | F(1,8) = 35.571 | <0.001 | 0.816 |  |
|  |  | Response x Drop type | F(1,8) = 1.455 | 0.262 | 0.154 |  |
|  |  | Bonferroni (Active vs. Random) | Mean Difference (Group 1 - Group 2) | Std. Error | p-value |  |
|  |  | Response | -0.075 | 0.105 | 0.497 |  |
|  |  | No-response | -0.326 | 0.114 | 0.021 |  |
| | Both T & E (T∩E) | RM-ANOVA (Response x Drop type) | F-value | p-value | Partial $\eta^2$ | 2B.3, bottom right |
|  |  | Response (within-subjects) | F(1,8) = 12.586 | 0.008 | 0.611 |  |
|  |  | Drop type (within-subjects) | F(1,8) = 4.787 | 0.060 | 0.374 |  |
|  |  | Response x Drop type | F(1,8) = 9.020 | 0.017 | 0.530 |  |
|  |  | Bonferroni (Active vs. Random) | Mean Difference (Group 1 - Group 2) | Std. Error | p-value |  |
|  |  | Response | -0.254 | 0.095 | 0.029 |  |
|  |  | No-response | 0.044 | 0.021 | 0.075 |  |

**Table S6.** Detailed statistical outputs for trial-wise decoder accuracy during reinstatement.

| Measure | Factors in analysis | Statistical output |  |  | Figure |
| --- | --- | --- | --- | --- | --- |
| Decoder accuracy on Reinstatement trials ($n = 9$) | RM-ANOVA (Drop type x Population) | F-value | p-value | Partial $\eta^2$ | 3B.2, left |
|  | Population (within-subjects) | F(1.636,13.091) = 3.848 | 0.006 | 0.325 |  |
|  | Drop type (within-subjects) | F(1,8) = 2.704 | 0.139 | 0.253 |  |
|  | Drop type x Population | F(2.040,16.317) = 2.872 | 0.085 | 0.264 |  |
|  | Bonferroni (Active vs. Random) | Mean Difference (Group 1 - Group 2) | Std. Error | p-value |  |
|  | All training (T) | -0.071 | 0.033 | 0.061 |  |
|  | All extinction (E) | -0.064 | 0.030 | 0.069 |  |
|  | All (T∪E) | -0.035 | 0.050 | 0.506 |  |
|  | Overlap (T∩E) | -0.097 | 0.029 | 0.011 |  |
|  | Only Training (T-E) | 0.030 | 0.024 | 0.248 |  |
|  | Only Extinction (E-T) | 0.006 | 0.038 | 0.888 |  |

**Table S7.** Detailed statistical outputs for trial-wise decoder accuracy during reinstatement following exclusion of specific active neuron sub-populations.

| Measure | Active neuron population | Factors in analysis | Statistical output |  |  | Figure |
| --- | --- | --- | --- | --- | --- | --- |
| Decoder accuracy on Reinstatement trials ($n = 9$) | All training (T) | RM-ANOVA (Response x Drop type) | F-value | $p$-value | Partial $\eta^2$ | 3B.3, top left |
| | | Response (within-subjects) | $F(1,8) = 16.358$ | 0.004 | 0.672 | |
| | | Drop type (within-subjects) | $F(1,8) = 3.726$ | 0.090 | 0.032 | |
| | | Response x Drop type | $F(1,8) = 5.228$ | 0.052 | 0.395 | |
| | | Bonferroni (Active vs. Random) | Mean Difference (Group 1 - Group 2) | Std. Error | $p$-value | |
|  |  | Response | -0.152 | 0.069 | 0.058 |  |
|  |  | No-response | 0.024 | 0.020 | 0.267 |  |
| | All extinction (E) | RM-ANOVA (Response x Drop type) | F-value | $p$-value | Partial $\eta^2$ | 3B.3, bottom left |
| | | Response (within-subjects) | $F(1,8) = 0.227$ | 0.646 | 0.028 | |
| | | Drop type (within-subjects) | $F(1,8) = 9.482$ | 0.015 | 0.542 | |
| | | Response x Drop type | $F(1,8) = 2.830$ | 0.131 | 0.261 | |
| | | Bonferroni (Active vs. Random) | Mean Difference (Group 1 - Group 2) | Std. Error | $p$-value | |
|  |  | Response | 0.010 | 0.067 | 0.881 |  |
|  |  | No-response | -0.272 | 0.115 | 0.045 |  |
| | Only Training (T-E) | RM-ANOVA (Response x Drop type) | F-value | $p$-value | Partial $\eta^2$ | 3B.3, top center |
| | | Response (within-subjects) | $F(1,8) = 23.942$ | 0.001 | 0.750 | |
| | | Drop type (within-subjects) | $F(1,8) = 0.523$ | 0.490 | 0.061 | |
| | | Response x Drop type | $F(1,8) = 2.625$ | 0.144 | 0.247 | |
| | | Bonferroni (Active vs. Random) | Mean Difference (Group 1 - Group 2) | Std. Error | $p$-value | |
|  |  | Response | 0.058 | 0.037 | 0.161 |  |
|  |  | No-response | -0.042 | 0.027 | 0.157 |  |
| | Only Extinction (E-T) | RM-ANOVA (Response x Drop type) | F-value | $p$-value | Partial $\eta^2$ | 3B.3, bottom center |
| | | Response (within-subjects) | $F(1,8) = 0.346$ | 0.573 | 0.041 | |
| | | Drop type (within-subjects) | $F(1,8) = 2.871$ | 0.129 | 0.264 | |
| | | Response x Drop type | $F(1,8) = 93.351$ | < 0.001 | 0.921 | |
| | | Bonferroni (*Active* vs. Random) | Mean Difference (Group 1 - Group 2) | Std. Error | $p$-value | |
|  |  | Response | 0.229 | 0.036 | < 0.001 |  |
|  |  | No-response | -0.408 | 0.080 | < 0.001 |  |
| | All (T∪E) | RM-ANOVA (Response x Drop type) | F-value | $p$-value | Partial $\eta^2$ | 3B.3, top right |
| | | Response (within-subjects) | $F(1,8) = 0.015$ | 0.904 | 0.002 | |
| | | Drop type (within-subjects) | $F(1,8) = 13.156$ | 0.007 | 0.622 | |
| | | Response x Drop type | $F(1,8) = 3.704$ | 0.090 | 0.316 | |
| | | Bonferroni (*Active* vs. Random) | Mean Difference (Group 1 - Group 2) | Std. Error | $p$-value | |
|  |  | Response | 0.056 | 0.097 | 0.579 |  |
|  |  | No-response | -0.341 | 0.122 | 0.024 |  |
| | Overlap (T∩E) | RM-ANOVA (Response x Drop type) | F-value | $p$-value | Partial $\eta^2$ | 3B.3, bottom right |
| | | Response (within-subjects) | $F(1,8) = 20.509$ | 0.002 | 0.719 | |
| | | Drop type (within-subjects) | $F(1,8) = 7.443$ | 0.026 | 0.482 | |
| | | Response x Drop type | $F(1,8) = 10.263$ | 0.013 | 0.562 | |
| | | Bonferroni (*Active* vs. Random) | Mean Difference (Group 1 - Group 2) | Std. Error | $p$-value | |
|  |  | Response | -0.182 | 0.057 | 0.012 |  |
|  |  | No-response | 0.036 | 0.024 | 0.165 |  |

